# A glucosylcholesterol-cytoskeleton axis links GBA2 loss-of-function to synaptic and mitochondrial pathology in Hereditary Spastic Paraplegia

**DOI:** 10.64898/2026.08.26.747028

**Authors:** A. Casotto, C. Sinisgalli, F. Terrin, L. Presicce, N. Facchinello, N. He, S. Marcotti, M. Dal Maschio, F. M. Santorelli, L. Laraia, L. Dalla Valle, N. Plotegher

## Abstract

**Background:** GBA2-associated hereditary spastic paraplegia (SPG46) is a rare autosomal recessive neurodegenerative disorder caused by loss-of-function mutations in *GBA2*, encoding the non-lysosomal glucocerebrosidase 2. GBA2 deficiency leads to glucosylceramide (GlcCer) accumulation and glucosylated cholesterol (GlcChol) depletion, causing cytoskeletal defects in immature neurons. However, the mechanisms linking lipid dysregulation to neuronal dysfunction remain poorly understood.

**Methods:** We modelled GBA2 loss of function by chronic pharmacological inhibition in mouse cerebellar granule neurons (CGNs) and assessed neuronal morphology, synaptic organization, Ca^2+^ dynamics, mitochondrial function and actin cytoskeleton during maturation. Proteomic profiling was performed in GBA2-inhibited and GlcChol-supplemented neurons. Findings were validated in a zebrafish *gba2* crispant model by evaluating motor behavior, cerebellar development, neuronal organization and mitochondrial function, and in patient-derived fibroblasts carrying a homozygous pathogenic *GBA2* variant (NM_020944). The role of RAC1 was studied in both neurons and patients’ cultured skin fibroblasts, and upon *rac1* pharmacological inhibition in zebrafish crispants.

**Results:** Chronic GBA2 inhibition impaired axonal outgrowth in immature CGNs but not neurite complexity in mature neurons, suggesting morphological compensation. Nevertheless, mature neurons displayed enlarged presynaptic terminals, impaired synaptic vesicle clustering and altered Ca^2+^ responses to potassium and glutamate, the latter associated with NMDA receptor redistribution without changes in total receptor levels. Mitochondrial alterations were observed in CGNs, patient fibroblasts and zebrafish, consistent with defective architecture of the mitochondrial network. Proteomics revealed convergent alterations in actin cytoskeleton, synaptic pathways and cellular metabolism following both GBA2 inhibition and GlcChol supplementation. GlcChol bidirectionally regulated RAC1 function, likely altering its spatial distribution rather than its global activation. Confocal imaging confirmed abnormal RAC1 and F-actin localization in patient fibroblasts. Zebrafish *gba2* crispants recapitulated motor deficits, Purkinje cell loss, motor neuron disorganization and mitochondrial abnormalities. Pharmacological Rac1 inhibition rescued motor behavior and neuronal organization, linking cytoskeletal disorganization to the observed phenotype in the zebrafish model.

**Conclusions:** Our findings identify a pathogenic GlcChol-RAC1-actin signalling axis linking lipid imbalance to synaptic disorganization, NMDA receptor redistribution and mitochondrial dysfunction in SPG46. The selective vulnerability of corticospinal neurons, cerebellar granule neurons and Purkinje cells may reflect their dependence on this pathway. Rac1 inhibition rescues disease phenotypes *in vivo*, highlighting this pathway as a promising therapeutic target.

## Background

*GBA2* encodes the non-lysosomal glucocerebrosidase 2 (GBA2), an enzyme localized at the cytoplasmic side of the ER and Golgi membranes [1]. GBA2 hydrolyzes glucosylceramides (GlcCer) to glucose and ceramides, and transglucosylates cholesterol to form β-D-glucosyl cholesterol (GlcChol) [2]. The related *GBA1* gene encodes the lysosomal glucocerebrosidase, which acts on the same substrates of *GBA2*, but within the lysosomal compartment. While biallelic pathogenic variants in *GBA1* cause Gaucher Disease (GD) [3], the most common lysosomal storage disorder, loss-of-function biallelic missense and nonsense mutations in *GBA2* cause a rare inherited form of Hereditary Spastic Paraplegia (HSP) type 46 [4,5]. HSP comprises a group of genetically diverse neurodegenerative disorders characterized primarily by progressive spasticity and weakness of the lower limbs. With over 90 genetic types identified, HSPs show significant clinical and molecular heterogeneity, since they are associated with impaired regulation of different functions including membrane/organelle trafficking, axon transport, mitochondrial activity and myelination process [6]. Despite its debilitating impact, there are currently no disease-modifying treatments, underscoring the urgent need for novel therapeutic strategies in HSPs [7,8]. *GBA2*-associated HSP (SPG46) is a complex rare form characterized by early onset, slow progression and a broad spectrum of neurological and systemic manifestations [4,5]. Patients typically present spastic gait, cerebellar ataxia, cognitive decline, peripheral neuropathy, scoliosis and cataract. Notably, cerebellar atrophy is a consistent and prominent feature of *GBA2*-HSP, distinguishing it from pure forms of HSP and suggesting selective vulnerability of cerebellar circuitry to GBA2 dysfunction. This is consistent with the high expression of GBA2 in cerebellar neurons ([9]; https://gtexportal.org/home/gene/GBA2) and with the well-established sensitivity of these neuronal populations to perturbations in lipid metabolism [10,11]. GlcCer accumulation, together with its metabolite glucosylsphingosine, has long been recognized as pathologically relevant in GD, as a consequence of defects in *GBA1* [12]. On the other hand, recent studies suggest that also GlcChol may be important in cell signaling and play a relevant role in neuronal function and in the aetiology of different diseases, including neurodegenerative disorders [2,13,14], similarly to the plant-derived glucosylated-sitosterol [15,16]. Loss-of-function mutations in *GBA2,* therefore, result in a dual lipid imbalance, GlcCer accumulation and GlcChol depletion, whose downstream consequences on neuronal function remain unclear [9]. GBA2 deficiency has been shown to impair cytoskeleton remodeling in mouse fibroblasts, likely by modulating actin polymerization and microtubule organization, and to cause defective neurite outgrowth in immature neurons [17,18]. These defects correlate with hyperactivation of RAC1, a small Rho GTPase that is a central regulator of actin cytoskeleton dynamics, cell polarity and membrane remodeling [19]. The pathological relevance of RAC1 dysregulation in the central nervous system is further underlined by the fact that *de novo* variants in *RAC1* itself cause a neurodevelopmental syndrome characterized by intellectual disability, brain malformations, spasticity and seldom cerebellar abnormalities [20,21]. Both gain-of-function and loss-of-function pathological variants in *RAC1* have been identified in patients, highlighting that tight control of RAC1 activity is essential for proper neuronal development and function. The activity of RAC1 is known to be modulated by the organization of membrane lipids and lipid rafts [22], suggesting that lipid alterations that are consequences of GBA2 deficiency may represent a direct upstream mechanism driving RAC1 dysregulation. Despite these advances, several critical issues remain unresolved. First, the respective contributions of GlcCer accumulation and GlcChol depletion to the observed neuronal defects have not been dissected, leaving open the question of which lipid species is the primary pathogenic driver. Second, while cytoskeletal defects have been characterized in immature neurons [18], the consequences of GBA2 loss on neuronal development, synaptic organization and neurotransmitter signaling in mature neurons remain still unknown. Finally, whether the RAC1-cytoskeleton axis represents a pharmacologically actionable target in this disease context has not been tested in cells or in a physiologically relevant *in vivo* model.

Here, we combine cellular, proteomic and imaging approaches to dissect the pathogenic consequences of GBA2 loss of function, from early morphological defects in immature cerebellar granule neurons (CGNs) to synaptic and functional alterations in mature neurons. We identify GlcChol depletion as a key upstream event linking GBA2 dysfunction to dysregulation of cytoskeletal and synaptic structures, to mitochondrial alterations and to Ca^2+^ signaling defects, the latter being mechanistically distinct from those observed in other lipid dyshomeostasis models [23]. These findings are validated through a *gba2* crispant zebrafish model and patient-derived fibroblasts carrying the homozygous p.Cys688Thr mutation [4], supporting their translational relevance. Across all models, RAC1 emerges as a central mediator of cytoskeletal dysregulation downstream of alterations in GlcChol levels. Moreover, RAC1 pharmacological inhibition rescues neuronal disorganization and motor deficits *in vivo*, identifying the GlcChol-RAC1-actin axis as a potential therapeutic target in GBA2-associated HSP.

## Methods

### Mice and zebrafish husbandry

C57BL/6 mice were maintained within the Animal Facility of the Department of Biology, at the University of Padova. Dark/light cycles were 12 h/12 h, temperature was kept between 21 and 24 °C and humidity between 60% and 65%. Wild type (WT) and transgenic zebrafish were maintained at the Zebrafish Facility of the University of Padova under standard guidelines [24]. Embryos were obtained by natural mating and raised at 28.5°C in Petri dishes containing Fish Water (50X: 25 g Instant Ocean (Aquarium systems, SS15-10), 39.25 g CaSO_4_ and 5 g NaHCO_3_ for 1 l), with a photoperiod of dark/light cycles of 12h/12h. The transgenic zebrafish lines Tg(mnx1:mGFP) [25] and Tg(aldoca:epNTR-TagRFP)^nub125^ [26] were used in this work.

### Primary mouse cerebellar granule neuron preparation, treatments and transfection

Primary cerebellar granule neurons (CGNs) were derived from C57BL/6 wild type postnatal mice (P5-P8). The brain was isolated and kept in dissection medium (15 nM Albumin (Roth), 10 mM Glucose (Sigma) in Dulbecco’s Phosphate Buffer Saline (PBS, Biowest)). Cerebella were isolated and incubated in Trypsin solution (1 mL/cerebellum, 0.1% Trypsin (Gibco), 1 mg/µL DNAse (Roche), 15 nM Albumin (Roth), 10 mM Glucose (Sigma) in PBS) at 37°C for 25 minutes. After enzymatic digestion, the supernatant was discarded and replaced with 4 mL of stop solution (30 nM Trypsin Inhibitor (Sigma), 1 mg/µL DNAsi (Roche), 15 nM Albumin (Roth), 10 mM Glucose (Sigma) in PBS). Cerebella were mechanically dissociated by gentle trituration and the resulting cell suspension was filtered through a 40-µm cell strainer (Sarstedt). Cells were resuspended in 10mL of Dulbecco’s Modified Eagle Medium (DMEM, Biowest) and pelleted. After removing the supernatant, cerebellar neurons were resuspended in Neurobasal medium (Gibco) supplemented with 2% v/v B27 (Gibco), 3% Penicillin-Streptomycin (Biowest) and 2.5% *L*-glutamine (Gibco). Cells were plated on poly-L-lysine-coated plates and incubated at 37°C with 5% CO_2_, replacing half of the culture medium with fresh medium every 3-4 days.

To inhibit GBA2 activity, neurons were incubated with the GBA2 inhibitor AMP-deoxynojirimycin (AMP-DNM; Cayman Chemical) at a final concentration of 1 nM from plating (DIV0) until 3, 9, or 14 days *in vitro* (DIV), depending on the experimental design. Control cells (CTRL) were treated with vehicle (0.1% DMSO). In selected experiments, β-glucosyl-cholesterol (10 μM, GlcChol, Sigma) was used to treat control neurons at DIV 13 for 24 hours. Chronic treatments were refreshed at each medium change.

When needed, 3-4 days after seeding, neurons were transiently transfected with pEGFP-N1 plasmid obtained from Clontech (Mountain View, CA, USA) and/or PSD95 expressing plasmid [27]. Lipofectamine 2000 (Invitrogen) was used as a transfection reagent and neurons were incubated with DNA:Lipofectamine (1:2) in OPTIMEM (Life Technologies) for 45 minutes, before the media was changed. Human fibroblasts from patients healthy (CTRL) and affected by hereditary spastic paraplegia (HSP) carrying the c.2063G>A (p.Cys688Thr) mutation (C688T) were cultured in DMEM, supplemented with fetal bovine serum (FBS, 10% Sigma Aldrich), 100 U/mL penicillin (Sigma Aldrich) and 100 μg/mL streptomycin (Sigma Aldrich) at 37 °C and 5% CO_2_.

### Western blot analysis

CGNs (DMSO, AMP-DNM and GlcChol) and patient-derived fibroblasts (CTRL and C688T) were seeded on 6-well plates (1.5x106 cells/mL and 5x105 cells/well, respectively) and lysed in RIPA buffer (20 mM Tris-HCl pH 7.5, 150 mM NaCl, 1 mM EDTA, 2.5 mM sodium pyrophosphate, 1 mM β-glycerophosphate, 1 mM sodium orthovanadate) supplemented with 1% protease inhibitor cocktail (Roche). After 30 minutes of ice incubation, samples were centrifugated for 30 minutes at 4°C and protein concentration in the supernatant was determined through the Pierce® BCA Protein Assay Kit following manufacturer’s instructions (Thermo Scientific) using an absorbance microplate reader (Victor X3, Perkin Elmer). Bovine serum albumin (BSA) was used as standard. Protein samples were denatured in 4X Sample Buffer (200 mM TrisHCl pH 6.8, 8% SDS, 400 mM DTT, 40% glycerol, 0.4% Bromophenol Blue (Sigma-Aldrich)) to reach a final 1X concentration. Then protein samples were resolved by electrophoresis on 8% tris-glycine polyacrylamide gel or gradient gel 4–20% Tris-MOPS-SDS gels (GenScript) and then transferred to polyvinylidenedifluoride (PVDF) membranes using a semi-dry Biorad transfer machine (TransBlot® Turbo TM Transfer System) with Transfer Buffer (BioRad). Membranes were blocked in saline Tris-buffered supplemented with 0.1% Tween® 20 (TBS-T: 20 mM Tris-HCl pH 7.4; 150 mM NaCl; 0.1% v/v Tween® 20) and 5% (w/v) non-fat dry milk for 1 h and then incubated overnight at 4°C with primary antibodies diluted in TBS-T plus 5% non-fat dry milk. The following primary antibodies were used: rat anti-GBA2 (1:20, gifted by Dagmar Wachten (23250757)), rabbit anti-GBA1 (1:1000, G4171 Sigma-Aldrich), mouse anti-HSP90 (1:2000, BD Biosciences), mouse anti-VGLUT1 (1:1000, Synaptic Systems), rabbit anti-DREBRIN (1:1000, MA1-20377 Invitrogen), mouse anti-transgelin-3 (1:100, sc-53932 Santa Cruz) rabbit anti-NMDAR2B (1:1000 AGC-003 Alomone labs), rabbit anti-pPAK1/2/3 (1:1000, SAB4503964 Sigma Aldrich), rabbit anti-RAC1 (1:1000, GTX100761 GeneTex), Phospho-Cofilin (Ser3) (77G2) (1:1000, 3313S Cell signaling), mouse anti-*β*-actin (1:1000, A3853 Sigma Aldrich), rabbit anti-TOM20 (1:2000, 11802-1-AP Proteintech), rabbit anti-MFN1 (1:1000, 13798 Proteintech), mouse anti-MFN2 (1:1000 H00009927 Abnova), rabbit anti-OPA1 (1:1000, Ab42364 Abcam). After washes, membranes were incubated for 1 hour at room temperature with Horse Radish-Peroxidase (HRP)-conjugated secondary antibodies diluted in TBS-T plus 5% non-fat dry milk: goat anti-mouse-HRP (1:80000, Sigma-Aldrich), goat anti-rabbit-HRP (1:16000, A9169 Sigma-Aldrich), goat anti-rat-HRP (1:2000, Sigma-Aldrich) diluted in 5% w/v skimmed milk in TBS-T. Membranes were visualized using Immobilon® Western HRP Substrate (Millipore) by VWR^®^ Imager Chemi Premium detector (VWR). ImageJ software was used to carry out the densitometric analysis of the detected bands.

### Glucocerebrosidases activity in cell lysates

Primary CGNs (DMSO and AMP-DNM) and fibroblast from patients (CTRL and C688T) cultured on 6-well plates (1.5x10^6^ cells/mL and 5x10^5^ cells/well, respectively) were lysed at the appropriate time point and confluence in hypotonic buffer (10 mM Hepes, 0.5 mM EDTA, pH 7.4) supplemented with 1% protease inhibitor cocktail (Roche) and homogenized at 4°C by pestle [1]. The total lysates were centrifuged and protein concentration in the supernatant was determined with Pierce® BCA Protein Assay Kit. According to the quantification, samples were diluted in PBS (20 μL, 0.1 μg/μL) and incubated for 90 minutes at 37°C in the dark with 40 μL of the substrate 4-methylumbelliferyl-b-D-glucopyranoside (4-MU, 3 mM, Sigma-Aldrich) in citrate phosphate buffer (Citric Acid 0.1 M; Na_2_HPO_4_ 0.2 M (Sigma-Aldrich)) at pH 4.5 supplemented with 0.2% taurodoexycholate (Sigma-Aldrich) for GBA1 assay, and at pH 6.0 for GBA2 assay. Afterwards, 240 μL of stop solution (Glicine 0.2 M at pH 10.4) at 4°C were added in each well and the fluorescence (λ_ex_ 360nm and λ_em_ 446nm) was measured in a fluorescence microplate reader (Victor X3, Perkin Elmer).

### RNA isolation and quantitative real-time PCR (RT-qPCR)

Total RNA was isolated from CGNs (DMSO and AMP-DNM) at 9 DIV using the RNeasy Mini Kit (Qiagen) following manufacturer’s instructions. Zebrafish total RNA was instead extracted from pools of 15-20 embryos, 10-12 larvae or single adult organs with TRIzol reagent (Thermo Fisher Scientific), according to the manufacturer’s instructions. RNA samples were treated with DNaseI (Promega). Complementary DNA (cDNA) was obtained from total RNA using iScript cDNA Synthesis Kit (Bio Rad) for cell samples, while High-Capacity cDNA Reverse Transcription Kit (ThermoFisher) was used for zebrafish samples. Real-time qPCR was carried out exploiting SensiFAST SYBR Lo-ROX Kit (Bioline) on a QuantStudio™ 6&7 Flex (Software 1.3 version, ThermoFisher) for CGNs samples, and SYBR Green PCR Master Mix (ThermoFisher Scientific) on a CFX384 (BioRad) for zebrafish cDNA. All samples were loaded in triplicates and transcripts levels were normalized for the appropriate housekeeping gene relative abundance. Data were analyzed applying the ΔΔCt algorithm. Primers used in this work are listed in **Additional files 2 - Table 1**.

### Mass Spectrometry to evaluate GlcChol level

CGNs were seeded at a density of 1 x 10^7^ cells per 10-cm dish. At DIV14, cells from two dishes were pooled and pelleted. For each sample, the entire pellet was homogenized and Cholesterol-D6 was added to each sample, as an internal standard. Lipids were extracted using a liquid-liquid extraction method according to Folch’s protocol [23]. The resulting clear solutions were analysed by Lipotype GmbH using an Agilent 1290 Infinity II system coupled to an Agilent 6495 triple quadrupole mass spectrometer Agilent 6495/1290 HPLC/MS/MS equipped with a 50 mm reversed-phase column as stationary phase. The mobile phase consisted of Ammonium formate 5 mM in Methanol/i-Propanol/Water. The detection was provided in Multi-Reaction-Mode (MRM) with two transitions each. Based on these data, limits of quantification (LOQ) and coefficients of variation (cv) for each lipid class were provided by Lipotype GmbH. The LOQ values were in the lower pg range, depending on the analyte. The average coefficient of variation for a complete set of analytes is <15. The amount of GlcChol was expressed in pmol per sample.

### Immunocytochemistry and confocal microscopy analysis

Primary CGNs (DMSO, AMP-DNM and GlcChol) and fibroblasts from patients were cultured onto glass coverslips in a 24-well plate (250.000 cells/well and 50.000 cells/well, respectively) and fixed for 20 minutes at room temperature using paraformaldehyde (4 % PFA, Sigma-Aldrich) dissolved in PBS. Cells were then permeabilized with PBS with 0.1% Triton for 20 minutes and then blocked with blocking solution (PBS supplemented with 5% Fetal Bovine Serum (FBS, Corning)) for 1 h at room temperature. Cells were incubated overnight at 4°C with the following primary antibodies: chicken anti-β-III-Tubulin (1:300 302 306 Synaptic System), mouse anti-VGLUT1 (1:300, 135 011 Synaptic Systems), rabbit anti-TAU (1:300, GTX112981 GeneTex), rabbit anti-NMDAR2B (1:300, AGC-003 Alomone labs), mouse-phalloidin (1:1000, ab176759 Abcam), rat anti-GBA2 (1:20, gifted by Dagmar Wachten (Korschen et al, 2012)), rabbit anti-RAC1 (1:1000, GTX100761 GeneTex). The day after, cells were washed with PBS and incubated for 1 hour at room temperature with the following secondary antibody (1:200 in blocking solution): goat anti-mouse Alexa Fluor 488 (A11029 Invitrogen), goat anti-mouse Alexa Fluor 568 (A11004 Invitrogen), goat anti-mouse Alexa Fluor 633 (A21052 Invitrogen), goat anti-rabbit Alexa Fluor 568 (A11004 Invitrogen), goat anti-rabbit Alexa Fluor 488 (A-11034 Invitrogen), goat anti-chicken Alexa Fluor 647 (A32933 Invitrogen), goat anti-rat Alexa Fluor 568 (A11077 Invitrogen). Nuclei were stained with Hoechst 33258 pentahydrate (bis-benzimide) (1:10.000 in H2O, Invitrogen) for 5 minutes. Coverslips were mounted using Mowiol (Calbiochem) and stored at 4°C.

For morphological analysis, images were acquired at 8-bit resolution over 1024x1024 pixel, through Leica SP5 confocal microscope using a 40X oil immersion objective. Neurite complexity analysis was performed using the NeuronJ plugin in Fiji-ImageJ [27,28]. Neurite tracings were performed semi-automatically by manually defining the start and end points of each process, with the plugin automatically following the neurite morphology. Total neurite length was measured for each neuron. Neuronal complexity was assessed by using the Sholl Analysis plugin in Fiji-ImageJ [29], centered on the soma with a step size of 5 µm, until no further intersections were detected. Total branching was quantified using a custom Fiji-ImageJ macro. Images were thresholded using the Otsu method, converted to binary masks, dilated three times, and filtered to remove particles smaller than 20 pixels². Masks were then eroded three times and skeletonized. Branches were detected using the Ridge Detection plugin (line width=10 pixels; sigma=3.39; lower threshold=0; upper threshold=1.36; minimum line length=10 pixels) (https://zenodo.org/records/845874). For synapses analysis, NMDAR2B receptor localization and F-ACTIN staining, immunofluorescence z-stack images were obtained on Zeiss LSM700 laser scanning confocal microscopy by using a 63X oil immersion objective. Synaptic density analysis was performed using the SynapCountJ plugin in Fiji-ImageJ with default parameters [30].

### Transmission Electron Microscopy

Primary CGNs or patient derived fibroblasts were seeded in 24-well plates (2 x 10^5^ and 50.000 cells/well, respectively). CGNs at 14 DIV and fibroblasts at approximately 80% confluence were fixed in a fixative buffer (2.5% glutaraldehyde in 0.1 M sodium cacodylate buffer pH 7.4) for 90 minutes at 4°C. Then, samples were post-fixed using 1% osmium tetroxide and 1% potassium ferricyanide in 0.1 M sodium cacodylate buffer for 1h at 4°C. Cells were then dehydrated in a graded ethanol series and embedded in an epoxy resin (Sigma-Aldrich). Ultrathin sections (60-70 nm) were obtained with an Ultrotome V (LKB) ultramicrotome, counterstained with uranyl acetate and lead citrate and viewed with a Tecnai G2 (FEI) transmission electron microscope (TEM) operating at 100kV. Synaptic vesicle distances were analyzed by the PointDensitySyn plugin for ImageJ and PointDensitySyn Python program, as described elsewhere [31].

### Ca^2+^ imaging

For live imaging experiments, CGNs were seeded at a density of 0.2 x 10^5^ cells on µ-Slide 8 well^high^ ibiTreat (ibidi Gmbh). At 14 DIV, cells were incubated at 37°C with 1 µM Fura-2 (Invitrogen) in imaging buffer (150 mM NaCl, 4.25 mM KCl, 4 mM NaHCO_3_, 1.25 mM NaH_2_PO_4_, 1.2 mM MgCl_2_, 1.2 mM CaCl_2_, 10mM D-glucose, 10 mM HEPES at pH 7.4) for 30 minutes. Fura-2-loaded cells were visualized by a 40× ultraviolet-permeable objective (Olympus Biosystems GmbH, Planegg, Germany) on an inverted microscope (Zeiss Axiovert 100, Jena, Germany). A monochromator (polychrome V, TILL-Photonics) controlled by a custom-made software (Roboscope, developed by Catalin Dacian Ciubotaru) was used to alternate excitation wavelengths (340 nm and 380 nm, respectively, 200 ms exposure time at each wavelength). A neutral density filter, UVND 0.6 (Chroma, USA) was used in the excitation pathway. Fluorescence emission was collected at 500-530 nm. Images were acquired every second, by a PCO SensiCam QE camera (Kelheim, Germany) [32]. For live imaging, L-glutamate acid monosodium salt hydrate (10 µM, Merck) was added to the cells. In selected experiments, cells were pre-incubated with the NMDA receptor antagonist D-AP5 (50 µM, Cayman chemicals) or the AMPA receptor antagonist NBQX (25 µM, Cayman chemicals), prior to glutamate stimulation. In additional experiments, cells were stimulated with 50 mM KCl to induce depolarization. At the end of every experiment, ionomycin (2 µM, Cayman chemicals) was added as positive control. ROIs were defined for each cell body and the integrated density of the two wavelengths were measured using ImageJ. After background removal, ratios between the signal at 380 nm and 340 nm were calculated at each time point. The resulting traces were plotted and the peak height was calculated for each cell. Basal Ca^2+^ of unstimulated cells was evaluated measuring the average over the first three points of the baseline before the stimulation.

### Mitochondrial membrane potential analysis in live cells

CGNs were seeded onto ibidi slides (1x10^5^ cells/well) and treated with DMSO (0.1%, CTRL) or AMP-DNM (1 nM) as described before. To investigate membrane potential in basal condition at 14 DIV, medium was removed and cerebellar neurons were incubated with Tetramethylrhodamine-methyl-ester (TMRM) (150 nM, Invitrogen) in Imaging Buffer (150 mM NaCl, 4.25 mM KCl, 4 mM NaHCO_3_, 1.25 mM NaH_2_PO_4_, 1.2 mM MgCl_2_, 1.2 mM CaCl_2_, 10 mM D-Glucose, 10 mM Hepes dissolved in cell culture water, 300 μL/well) for 20 minutes. The mix was removed, and cells were incubated with DAPI (1:10000 in Imaging Buffer) for 5 minutes. After washes, fresh imaging buffer was added (300 μL/well) and membrane potential was evaluated with confocal microscope Leica SP5 under a 40X objective. Quantification of TMRM fluorescence intensities was performed by Fiji, setting a common threshold to all the images and quantifying the average fluorescence intensity per field of view. Fibroblasts from patients (CTRL and C688T) were cultured on 12-well plates (2x10^5^ cells/well) and incubated with TMRM (150 nM) and MitoTracker green^TM^ FM (1:1000, Thermofisher scientific) for 20 minutes. Cells were then harvested in PBS and fluorescence of both dyes were detected by BD LSRFortessa™ X-20 Cell Analyzer (BD Biosciences). Data were acquired on live single cells after exclusion of debris by forward and side scatter gating. TMRM fluorescence was detected in the PE-Texas Red channel, while MitoTracker™ Green fluorescence was collected in the FITC channel. At least 10,000 events per sample were recorded and analyzed using BD FACSDiva™ Software. Unstained cells were used as negative controls, while cells stained individually with either TMRM or MitoTracker green^TM^ were used to set detector voltages and define acquisition parameters.

### Rac1-GTP pull-down assay

CGNs were cultured into three different dishes (1x10^7^ cells/dish) and chronically treated with DMSO (0.1% CTRL), AMP-DNM (1 nM), or with GlcChol (10 μM) added on 13 DIV and kept for 24 hours. On 14 DIV, medium was removed and Rac1 pull-down assay was performed. Rac1 pull-down is a simple assay to monitor Rac1 GTPase activation. The kit of the assay kit provides a GST-fusion protein containing the p21-binding domain (PBD) of human p21-activated protein kinase 1 (PAK1), along with glutathione agarose resin, containing 0.05% sodium azide and functionalized to PAK1, to specifically pull down active Rac1. Active Rac1 (Rac1-GTP) binds to the PBD of PAK1, leading to its activation. Therefore, it can be used as a probe to specifically isolate the active form of Rac1. Briefly, culture medium was removed and cerebellar neurons were lysed in RIPA buffer (250 μL/dish) supplemented with protease inhibitor cocktail (Roche), using a Double Blade Cell Scraper (Alkali Scientific™). Protein concentration in clarified lysed samples was determined with Pierce® BCA Protein Assay Kit (Thermo Fischer Scientific), following the standard protocol. Glutathione-agarose resin (Millipore) was functionalized with GST or GST-human PAK1-PBD and equilibrated in PBS. Neuronal lysates (250 μL) were incubated with the resin at 4°C for 1 h under rotation. After incubation, samples were washed three times with PBS, and the bound proteins were subsequently eluted in sample buffer 2X by heating at 95°C in a thermoblock. Eluted samples (5 μL) and cell lysates (40 μL) were separated by 10% SDS-PAGE, probed with anti-RAC1 (1:1.000, GeneTex) and the respective goat anti-rabbit IgG-HRP (1:16.000, Sigma), and detected in order to measure active RAC1.

### Proteomic sample preparation

CGNs were seeded on 10 cm plates (1x10^7^ cells/dish). On 14 DIV, cells were washed with ice cold PBS twice and lysed by the addition of lysis buffer (0.4% v/v NP-40 in PBS) supplemented with 1% protease inhibitors (Roche). Lysis was finalized by four freeze-thaw cycles using liquid nitrogen. Cellular debris were removed by centrifugation at 20.000g for 30 minutes at 4 °C. After centrifugation, the protein concentration was adjusted to 0.87 mg/mL. For each sample, 75 µg of total proteins were further diluted in 100 mM Tetraethylammonium bromide (TEAB) and with 200 mM tris(2-carboxyethyl)phosphine (TCEP) for cysteine reduction for 1 h at 55 °C. 375 mM Iodoacetamide (IAA) was successively used for alkylation for 30 min at room temperature in the dark. After this step, proteins were precipitated overnight with 6 volumes of cold acetone at -20 °C. The following day the protein pellets were collected by centrifugation at 8000g at 4 °C for 10 minutes and resuspended in TEAB 100 mM before digestion. Protein digestion was performed by adding trypsin in a 1:80 enzyme:substrate ratio and left overnight at 37 °C, with gentle shaking in a ThermoMixer (Eppendorf®). After digestion, 75 µg of peptides were successively labelled with TMTpro™ 16plex reagents kits (Thermo Fisher). Experimental replicates were labelled with different TMT-batches concatenated by a pooled internal reference sample for each TMT-batch, which consisted of an additional sample obtained by pooling equal amounts of each sample from the same TMT-batch. TMT reaction was allowed for 2 hours and quenched with 5% hydroxylamine. All samples related to each batch were pooled and dried in SpeedVac (Eppendorf EP022822993) before desalting with Sep-Pak® Plus C18 cartridges. Peptides were eluted with 40% and 60% of acetonitrile in 0.1% of trifluoroacetic acid (TFA) and dried before injection of 30 µg in the UHPLC system (Dionex U3000) for high-pH fractionation. The separation of the peptides was carried out at a constant flowrate of 5 µl min^-1^ on a CSH C18 Acquity UPLC M-Class Peptide column, 130 Å, 1.7 µm, 300 µm × 150 mm (Waters, 186007563) using a 100 min linear gradient from 5 to 35% of mobile phase B (acetonitrile) with a subsequent 15 minutes gradient to 70%, before 5 min re-equilibration with 95% of mobile phase A (5mM ammonium bicarbonate, pH 10). 60 time-based fractions were pooled in 30 fractions in the collection plates. Clean-up of the fractions was performed by EvoTip according to the manufacturer’s instructions.

### LC-MS data collection

The EvoTips (EvoSep, EV2003) were loaded on the Evosep One module (EvoSep EV-1000) coupled to an Orbitrap Eclipse™ Tribid™ mass spectrometer (ThermoFisher Scientific). Peptides were loaded onto the EASY-Spray™ C18 column, 2 µm, 100 Å, 75 µm × 15 cm (Thermo Fisher Scientific, ES804) using the standard “30 samples per day” Evosep method. The method eluted the peptides with a 44 min gradient ranging from 5% to 90% acetonitrile with 0.1% formic acid. The MS acquisition was performed in data dependent-MS3 with real-time-search (RTS) and a FAIMS interface switching between CVs of −50 V and −70 V with cycle times of 2 s and 1.5 s, respectively. The data dependent acquisition mode was run in a MS1 scan range between 375 and 1500 *m/z* with a resolution of 120000, and a normalized gain control (AGC) *target* of 100%, with a maximum injection time of 50 ms. RF Lens set at 30%. Filtering of the precursors was performed using peptide monoisotopic peak selection (MIPS), including charge states from 2 to 7, dynamic exclusion of 120 s with ±10 ppm tolerance excluding isotopes, and a precursor fit of 70% in a window of 0.7 m/z with an intensity threshold of 5000. Selected precursors for further MS2 analysis were isolated with a window of 0.7 *m/z* in the quadrupole. The MS2 scan was performed over a range of 200-1400 *m/z*, collecting ions with a maximum injection time of 35 ms and normalized AGC target of 300% MS2 fragmentation was operated with normalized HCD collision energy at 30%. Fragmentation spectra were searched against the fasta files from the human Uniprot database (reviewed) in the RTS, set with tryptic digestion, TMT-11plex as fixed modification on Lysine (K) and N-Terminus together with cysteine (C) carbamidomethylation, and oxidation of methionine (M) as variable modification. 1 missed cleavage and 2 variable modifications were allowed with a maximum search time of 35 ms. FDR filtering was enabled with 1 as correlation and 5 ppm of precursor tolerance. Precursors identified via RTS in the MS2 scan were further isolated in the quadrupole with a 2 *m/z* window, maximum injection time of 86 ms and normalized AGC target of 300%. The further MS3 fragmentation was operated with a normalized HCD collision energy at 50% and fragments were scanned with a resolution of 50000 in the range of 100 to 500 *m/z*. The MS performances were monitored by quality control of an in-house standard of HeLa cell lysate, both at the beginning and the end of each sample set.

### LC-MS data analysis

Mass spectrometric raw files were analyzed with Proteome Discoverer 2.4 (Thermo Fisher Scientific) by using the built-in TMTpro Reporter ion standard quantification workflows. The search was run setting trypsin as enzyme (allowing maximum 2 missed cleavages), TMTpro16plex and carbamidomethylation of cysteine (C) as fixed modifications, while methionine (M) oxidation and acetylation of protein N-termini as variable modifications. Sequest search engine was used to match the MS2 spectra in the Uniprot homo sapiens database (Swiss-Prot reviewed including isoforms) with a precursor mass tolerance of 10 ppm and fragment mass tolerance of 0.6 Da. Percolator was used to score the results and to filter at 1% FDR. Reporter ion quantification was performed on MS2 spectra by applying isotopic error correction. Normalization and scaling were not included in the Proteome Discovery analysis and were performed successively on the protein result table.

Here, as a first step, contaminant proteins and proteins not identified as Master Protein or as Master Protein Candidate were removed. Moreover, all the proteins identified with a sum of Unique + Razor peptides below 2 were removed. Loading normalization was performed summing the intensities for each TMT channel and calculating the respective correction factor on the average of the summed intensities. Hence, each protein intensity was normalized for the respective channel correction factor. Normalized intensities were further considered to calculate the average among the replicates and the fold changes (FC) for compounds-treated samples over the control samples. Only proteins identified in all three replicates were used for further analysis. The obtained values were log2 transformed. A two-sided T-test was performed on the normalized data and obtained p-values were -log10 transformed. Volcano plots were obtained plotting the log2 Fold Change vs -log10 (p-value) in Prism 11 (GraphPad Software).

### Generation of zebrafish F0 knock-out (crispants) for the *gba2* gene

Genetic ablation of *gba*2 gene in zebrafish WT and transgenic larvae was obtained by CRISPR-Cas9 technology according to Masato and coworkers [33]. Three guide RNAs (gRNAs) designed to target the first three exons of the gene were used to achieve *gba2* silencing, maximizing mutagenesis efficiency. For gRNA assembly, each CRISPR RNA (crRNA) was annealed individually with an equal molar amount of trans-activating crRNA (tracrRNA; Alt-R™ CRISPR-Cas9 tracrRNA, 5 nmol; 1072532, IDT). crRNA-tracrRNA duplexes were prepared diluting the components in Duplex Buffer (IDT) to a final concentration of 57 μM and incubated at 95 °C for 5 minutes to allow annealing. For microinjection, each gRNA was used at a final concentration of 9.5 μM and combined with 10 μM Alt-R™ S.p. HiFi Cas9 Nuclease V3 (#1081060, IDT). gRNA-Cas9 ribonucleoprotein (RNP) complexes were incubated at 37°C for 5 minutes prior to injection. RNP complexes were injected into zebrafish embryos at the single-cell stage. The gRNAs targeting *gba2* were designed using the Breaking-Cas web tool and the IDT CRISPR design platform and are listed in **Additional files 2 - Table 2**. crRNAs for each target site were selected based on optimal predicted on-target efficiency and minimal off-target potential.

To verify the occurrence of possible side effects of gRNA injection, two scrambled gRNAs (Alt-R Cas9 Negative Control crRNA #1 (1072544, IDT) and #2 (1072545, IDT), 2 nmol) were adopted (see **Additional file 1, Fig. S4D**). Given the absence of side effects after injection, uninjected WT embryos were considered as controls.

### Genomic DNA extraction from larvae

To validate gRNA activity, genomic DNA was extracted from single larvae through the HotSHOT protocol [34] and was used to amplify the genomic regions spanning each CRISPR target site by PCR. PCRs were performed using HOT FIREPol^®^ Blend Master Mix (04-25-00115, Solis Biodyne) and the following program: 95°C for 12 min, 40 cycles of 95°C for 30 sec, 60°C for 20 sec, 72°C for 30 sec and a final elongation step at 72°C for 8 min. The three locus-specific primers are listed in **Additional files 2 - Table 3**.

### Morphological analysis of zebrafish larvae

Morphological traits, comprising standard body length (from snout to tail), eyes area, head length (from snout to posterior hindbrain), inter-ocular distance and swim bladder size, were analyzed in 5-dpf zebrafish larvae. After anaesthesia with 0.04% tricaine (MS-222, tricaine methanesulfonate, Sigma), larvae were arranged sideways or dorsally in 2% methyl-cellulose in H_2_O on microscope slides and photographed under a Leica M165 FC stereoscopic microscope equipped with a Leica DCF7000 T digital camera. Differences between crispants and control larvae were analysed with Fiji-ImageJ software (v. 2.14.0). Muscle birefringence index was analysed on 5-dpf larvae as previously described [35].

### Zebrafish larvae swimming behavioral analysis

Swimming behavioral analyses were performed as previously described [33], using a DanioVision™ automated tracking system (Noldus Information Technology). Briefly, 5-dpf zebrafish crispants and control larvae were singularly transferred to a 48-wells plate and subjected to a light-dark locomotor assay consisting of a pre-adaptation for 10 min in dark, followed by three cycles of 10 min in the light and 10 min in the dark. The tracking of the distance covered by larvae during the test was obtained through Ethovision XT software.

### Cerebellar fluorescence analysis in Tg(aldoca:epNTR-TagRFP)^nub125^ zebrafish larvae

Transgenic Tg(aldoca:epNTR-TagRFP)^nub125^ zebrafish larvae at one-cell stage were injected as previously described to obtain transgenic F0 *gba2* crispants. Starting from 30 hpf, zebrafish larvae were exposed to 1-phenyl-2-thiourea (PTU, Sigma Aldrich) to delay the onset of cutaneous pigmentation. PTU was renewed every 24h until 5 dpf. At 5 dpf, fluorescent transgenic *gba2* crispants and uninjected control larvae were anaesthetized with tricaine, mounted dorsally in low-melting 1% agarose and acquired with a Nikon C2 confocal microscope under a 20X objective (5 µm z-stack). For the fluorescence analysis, the sum of z-stacks projections was manually outlined and analyzed with Fiji-ImageJ for the mean fluorescence of *aldoca* positive cells. To evaluate cerebellar volume, images were transformed to 8-bit, filtered with Gaussian blur (Radius:1) and thresholded with Huang method. Then, Close and Fill Holes were applied to reconstitute the whole cerebellar structure. The 3D Object Counter plugin was used and the sum of total volumes was obtained. Data were normalized on the head volume approximated by multiplying head length (from olfactive bulbs to otic vesicle), head width (from one eye to the other), and head thickness (z-stacks number × thickness).

### Mitochondrial membrane potential analysis in zebrafish

For the analysis of mitochondrial membrane potential, zebrafish *gba2* crispants and control larvae at 4 dpf were incubated in the dark with TMRM (Thermo Fisher Scientific) at a final concentration of 300 nM dissolved directly in the water. After 24 hours, they were anaesthetized, mounted sideways in low-melting 1% agarose and acquired with a Nikon C2 confocal microscope under a 20X objective (7µm z-stack). Fluorescence analysis was performed on z-stack maximum projection on Fiji-ImageJ.

### Motor neurons image acquisition in Tg(mnx1:GFP) zebrafish larvae

Transgenic Tg(mnx1:mGFP) zebrafish larvae at one-cell stage were injected as previously described to obtain transgenic F0 *gba2* crispants. At 3 dpf, fluorescent transgenic *gba2* crispants and uninjected control larvae were anaesthetized with tricaine, mounted sideways in low-melting 1% agarose and imaged by z-stacks (2.5 µm z-stack) throughout their whole thickness with a Nikon C2 confocal microscope under a 20X objective.

### AFT-based image analysis

Image pre-processing of both fibroblasts and zebrafish larvae was performed in Fiji [36]. Each z-slice was saved as a separate image and segmented with Otsu automatic thresholding to obtain a binary mask. Three rounds of binary dilation/erosion were performed to fill holes and smooth edges. The signal was then skeletonized. For zebrafish confocal images, the contribution from the spinal cord was masked out with the “Analyze Particles” function, by keeping only objects with an area of less than 300 px. Subsequent alignment analysis was performed with AFT [36], an open-source tool to measure alignment of fibrillar features in bioimages. AFT parameters were chosen as follows: window size of 25 px, neighborhood radius of 3 vectors, and a filter on blank regions allowing to analyze only windows where the average signal was higher than 10/255 a.u. The results of the alignment analysis are displayed as wrapped color heatmaps, highlighting the local neuron angle.

### Rac1 inhibitor treatment in zebrafish

Treatment with RAC1 inhibitor NSC23766 (Cayman chemicals) was performed on *gba2* zebrafish crispants and uninjected controls by adding the compound directly in Fish water at a final concentration of 10 µM. After 24 h of incubation, larvae were rinsed in Fish water and processed for further rescue experiments as described previously.

### Statistical analyses

Statistical analyses were performed using GraphPad Prism. Data normality was assessed before analysis, and appropriate statistical tests were selected accordingly. Two-group comparisons were performed using Student’s t-test, whereas multiple-group comparisons were analyzed by one-way or two-way ANOVA with appropriate post hoc corrections. Outliers were excluded when required based on GraphPad Prism analysis. Results are shown as mean ± SEM, with p < 0.05 considered statistically significant.

## Results

### Chronic GBA2 inhibition induces biochemical remodeling without sustained morphological defects

To model the GBA2 loss of function that characterizes *GBA2* mutations associated with HSP, cerebellar granule neurons (CGNs) were chronically treated with AMP-Deoxynojirimycin (AMP-DNM, 1 nM), a potent GBA2 inhibitor [18]. The cerebellar involvement in SPG46 disease form provides a strong rationale for the use of CGNs as a mammalian cellular model to investigate the pathogenic mechanisms downstream of *GBA2* loss of function. Biochemical alterations were first analyzed at early stages of maturation, i.e. at 3 and 9 days *in vitro* (DIV). Western blotting showed a significant 2.8-fold increase in GBA2 protein at 3 DIV, suggesting compensation for the inhibition by increased enzyme expression. GBA1 protein levels, evaluated to verify compensatory or off-target effects, displayed no significant changes (p value = 0.065) (**Fig. 1A**). At 9 DIV, both GBA2 and GBA1 protein levels were significantly increased (∼1.4-fold), suggesting that either their synthesis was increased or their degradation was hampered (**Fig. 1B**). RT-qPCR analysis revealed that *Gba2* exhibited a non-significant trend toward reduced transcripts expression (p=0.07), whereas *Gba1* mRNA remained unchanged, indicating no alterations in its transcriptional regulation (**Fig. 1C**). Enzymatic assays at 9 DIV revealed an 80% reduction of GBA2 activity as compared to the control, while GBA1 activity was unaffected despite elevated protein levels (**Fig. 1D**). Confocal imaging at 9 DIV further confirmed GBA2 presence in both control and treated conditions (**Fig. 1E**). These data indicated effective and specific GBA2 inhibition, accompanied by possible reduction in GBA2 degradation. Increase in GBA1 levels with unchanged overall enzymatic activity suggested the accumulation of deglycosylated immature forms of the protein [37].

**Figure 1.**
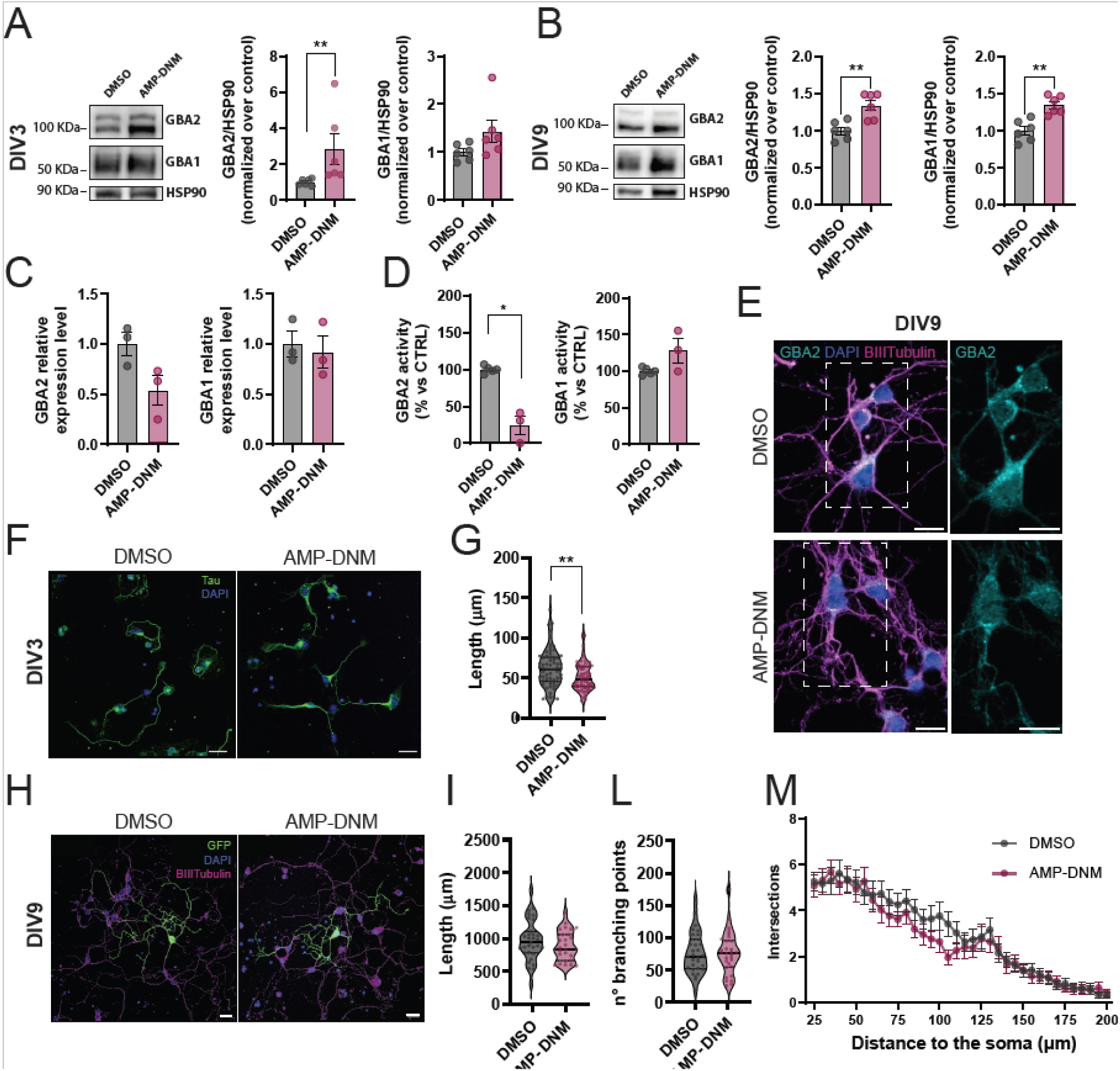
Neuronal development and morphology in GBA2-inhibited cerebellar granule neurons (CGNs) are altered. (A–B) GBA2 and GBA1 protein levels in CGNs treated with AMP-DNM (1 nM) at DIV3 (A) and DIV9 (B) (n = 6, 3 independent experiments; Mann-Whitney test and Welch’s t-test, respectively; A: GBA2 p = 0.0022, GBA1 p = 0.065; B: GBA2 p = 0.0033, GBA1 p = 0.0012). (C) RT-PCR analysis of GBA2 and GBA1 mRNA expression in CGNs at DIV9 treated with AMP-DNM (1 nM) (n = 3, 3 independent experiments; Welch’s t-test; GBA2 p = 0.0701, GBA1 p = 0.7). (D) GBA2 and GBA1 enzymatic activity in CGNs at DIV9 treated with AMP-DNM (1 nM) (n = 3, 3 independent experiments; Welch’s t-test; GBA2 p = 0.0022, GBA1 p = 0.22). (E) Representative immunofluorescence images of CGNs at DIV9 treated with AMP-DNM (1 nM), stained for GBA2 (cyan), β-III-Tubulin (magenta), and DAPI (blue). Scale bar, 10 μm. (F–G) Representative images (F) and quantification (G) of axon outgrowth in CGNs at DIV3 treated with AMP-DNM (1 nM), stained with anti-Tau (green) and DAPI (blue) (n ≥ 65, 3 independent experiments; Mann-Whitney test; p = 0.0033). Scale bar, 20 μm. (H–M) Representative images (H) and morphological quantification of GFP-transfected CGNs at DIV9 treated with AMP-DNM (1 nM), stained for β-III-Tubulin (magenta) and DAPI (blue): total neuronal length (I), branching point (L), and Sholl analysis (M) (n ≥ 29, 3 independent experiments; Welch’s t-test and two-way ANOVA; all p > 0.26). Scale bar, 20 μm. All data were tested for normality using the Shapiro-Wilk test.

We next assessed neuronal morphology of GBA2-inhibited CGNs as compared to DMSO-treated controls. Consistent with previous reports showing reduced neurite outgrowth in GBA2-inhibited neurons [18], axonal length was significantly reduced by about 20% at 3 DIV (**Fig. 1F, G**). However, at 9 DIV, detailed single-cell analysis of GFP-transfected neurons revealed no differences in total neurite length, branching, or overall neuronal complexity between GBA2-inhibited neurons and controls (**Fig. 1H, I, L, and M**) [27]. Since morphological defects were resolved over time, compensatory mechanisms may be in place to counteract the defective axonal outgrowth observed at earlier stages.

### Chronic GBA2 inhibition alters presynaptic ultrastructure and Ca^2+^ dynamics in mature neurons

To further assess if the reduced GBA2 function could affect CGNs neuronal activity at later stages, we grew CGNs until DIV 14, when functional synaptic structures are typically observed in cultured neurons. We hypothesized that any compensatory measures in place may not be able to level out all structural and functional defects in neurons at later stages. Thus, we examined synaptic organization exploiting confocal and transmission electron microscopy (TEM) (**Fig. 2A** and **2E**), as we had previously done in other models [27,33]. The density of glutamatergic excitatory synapses, measured as the colocalization between the pre-synaptic marker Vesicular Glutamate Transporter 1 (VGLUT1) and the post-synaptic marker Post Synaptic Density protein 95 (PSD-95), was unchanged between control and AMP-DNM-treated neurons (**Fig. 2B**). However, the number of VGLUT1 punctae, possibly marking pre-synpatic terminals, was increased by 33% in treated cultures compared to controls (**Fig. 2C**), which was confirmed by a significant ∼1.5-fold elevation of VGLUT1 protein level by western blot analysis (**Fig. 2D**). To investigate the ultrastructural features of the presynaptic terminals and the organization of synaptic vesicles pools, TEM imaging was performed (**Fig. 2E**). Image analysis revealed enlarged presynaptic terminals in treated neurons (50% increase of the area) compared to controls (**Fig. 2F**). Despite a non-significant difference in synaptic vesicle density (**Fig. 2G**), average inter-vesicle distance increased from 216.9 ± 0.7 nm in controls to 290.9 ± 0.4 nm following AMP-DNM treatment, while vesicle-to-PSD distance increased from 179.2 ± 2.8 nm to 263.5 ± 3.0 nm (**Fig. 2H, I**). These findings indicate expansion of the presynaptic compartment and increased vesicle dispersion, pointing towards a reduction in the pool of ready releasable vesicles.

**Figure 2.**
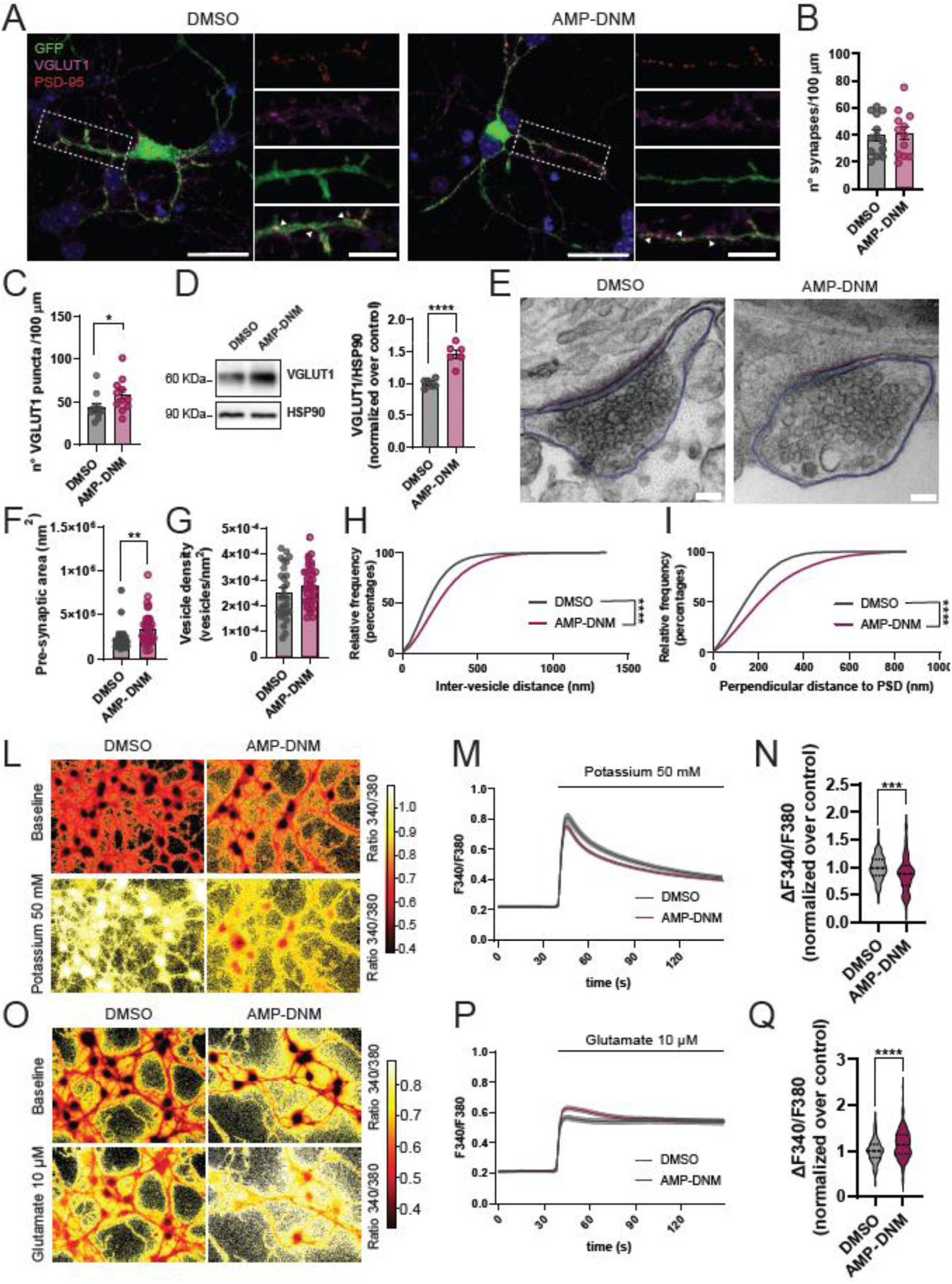
Chronic GBA2 inhibition alters presynaptic organization and Ca²⁺ dynamics in CGNs. (A) Representative confocal images of excitatory synapses in CGNs at DIV14 stained for the presynaptic marker VGLUT1 and the postsynaptic marker PSD-95 (scale bar, 10 µm; inset, 5 µm). Arrowheads indicate representative sites of VGLUT1/PSD-95 colocalization. (B) Quantification of synaptic density measured as VGLUT1/PSD-95 colocalization (n ≥ 12, 3 independent experiments; Welch’s t-test; p = 0.81). (C) Quantification of VGLUT1 puncta (n ≥ 12, 3 independent experiments; Mann–Whitney test; p = 0.034). (D) Western blot analysis and quantification of VGLUT1 protein levels in CGNs at DIV14 treated with AMP-DNM (1 nM) (n = 6, 3 independent experiments; Welch’s t-test; p = 0.0005). (E) Representative transmission electron microscopy (TEM) images of presynaptic terminals in control and AMP-DNM-treated CGNs at DIV14 (scale bar, 100 nm). The presynaptic terminal perimeter is outlined in blue and the postsynaptic density (PSD) in purple. (F) Quantification of presynaptic terminal area (n ≥ 27, 3 independent experiments; Mann–Whitney test; p = 0.001). (G) Quantification of synaptic vesicle density (n ≥ 27, 3 independent experiments, Welch’s t-test; p = 0.216). (H) Cumulative frequency distribution of vesicle-to-PSD distance (n ≥ 27, 3 independent experiments, Kolmogorov–Smirnov test; p < 0.0001). (I) Cumulative frequency distribution of inter-vesicle distance (n ≥ 27, 3 independent experiments, Kolmogorov–Smirnov test; p < 0.0001). (L) Representative ratiometric Fura-2 images (340/380 nm) of DMSO- and AMP-DNM-treated CGNs before and after stimulation (50 mM). (M) Fura-2 Ca²⁺ imaging traces (± SEM) in CGNs at DIV14 following KCl stimulation (50 mM). (N) Quantification of Ca²⁺ peak amplitude upon KCl stimulation (n ≥ 138, 3 independent experiments; Welch’s t-test; p = 0.0004). (O) Representative ratiometric Fura-2 images (340/380 nm) of DMSO- and AMP-DNM-treated CGNs before and after glutamate stimulation (10 µM). (P) Fura-2 Ca²⁺ imaging traces (± SEM) in CGNs at DIV14 following glutamate stimulation (10 µM). (Q) Quantification of Ca²⁺ peak amplitude upon glutamate stimulation (n ≥ 146, 3 independent experiments; Mann–Whitney test; p < 0.0001). All data were tested for normality using the Shapiro–Wilk test.

To determine whether these structural alterations affect neuronal function, we analyzed cytosolic Ca^2+^ dynamics using Fura-2 imaging by inducing neuronal depolarization with potassium chloride (KCl, 50 mM) or upon stimulation with non-toxic glutamate concentration (10 µM). While basal Ca^2+^ levels were unchanged (see **Additional file 1, Fig. S1**), potassium stimulation in GBA2-inhibited CGNs induced a lower peak of cytosolic Ca^2+^ response compared to controls (**Fig. 2L-N**). Conversely, stimulation with glutamate resulted in an increased cytosolic Ca^2+^ peak in treated CGNs (**Fig. 2O-Q**). Altogether, these data demonstrate that chronic GBA2 inhibition disrupts synaptic function, reflected in elevated VGLUT levels, impaired synaptic vesicle distribution, and dysregulated Ca^2+^ signaling. The diminished KCl response may suggest compromised presynaptic dynamics, such as impaired vesicle release or unbalanced vesicle pools, consistent with altered synaptic vesicles organization observed by TEM, but may be also coupled to glutamate receptor dysfunction [38,39]. It may also indicate impaired voltage-gated calcium channel (VGCC) function or localization. On the other hand, the increased glutamate-evoked signal may suggest enhanced activation of glutamate receptors, potentially due to changes in their abundance, conductance or distribution, or impaired Ca^2+^ handling, for example due to an impairment in the mitochondrial buffering capacity caused by mitochondrial impairment, as observed in *Gba1* knock-out mice primary neurons [23].

### Mitochondrial morphology and function are altered in GBA2-inhibited CGNs

The imbalance in stimulus-specific cytosolic Ca^2+^ responses suggested a potential impairment in intracellular Ca^2+^ handling; thus, we next assessed mitochondrial function, considering its pivotal role in Ca^2+^ buffering and in tuning neuronal Ca^2+^ homeostasis [40]. We evaluated mitochondrial membrane potential (ΔΨm) by live-cell confocal imaging using the dye tetramethylrhodamine methyl ester (TMRM). AMP-DNM treatment resulted in a significant increase in TMRM fluorescence compared to control CGNs (**Fig. 3A, B**). In parallel, analysis of mitochondrial morphology resulted in a significant increase of mitochondrial area, perimeter and Feret’s diameter in AMP-DNM-treated neurons, implying enlarged mitochondria (**Fig. 3C-F**). MFN1 and TOM20 showed reduced expression by Western blot, whereas MFN2 and OPA1 remained unchanged (**Fig. 3G-M**). When considered together with the enlarged mitochondrial morphology observed by TEM, these findings are consistent with changes in mitochondrial architecture, including a possible remodeling of the outer mitochondrial membrane [41], rather than with a generalized disruption of the mitochondrial fusion machinery. In this context, the increased TMRM signal may be indicative of functional changes within the mitochondrial network rather than mitochondrial hyperpolarization, possibly reflecting an early adaptive stress response.

**Figure 3.**
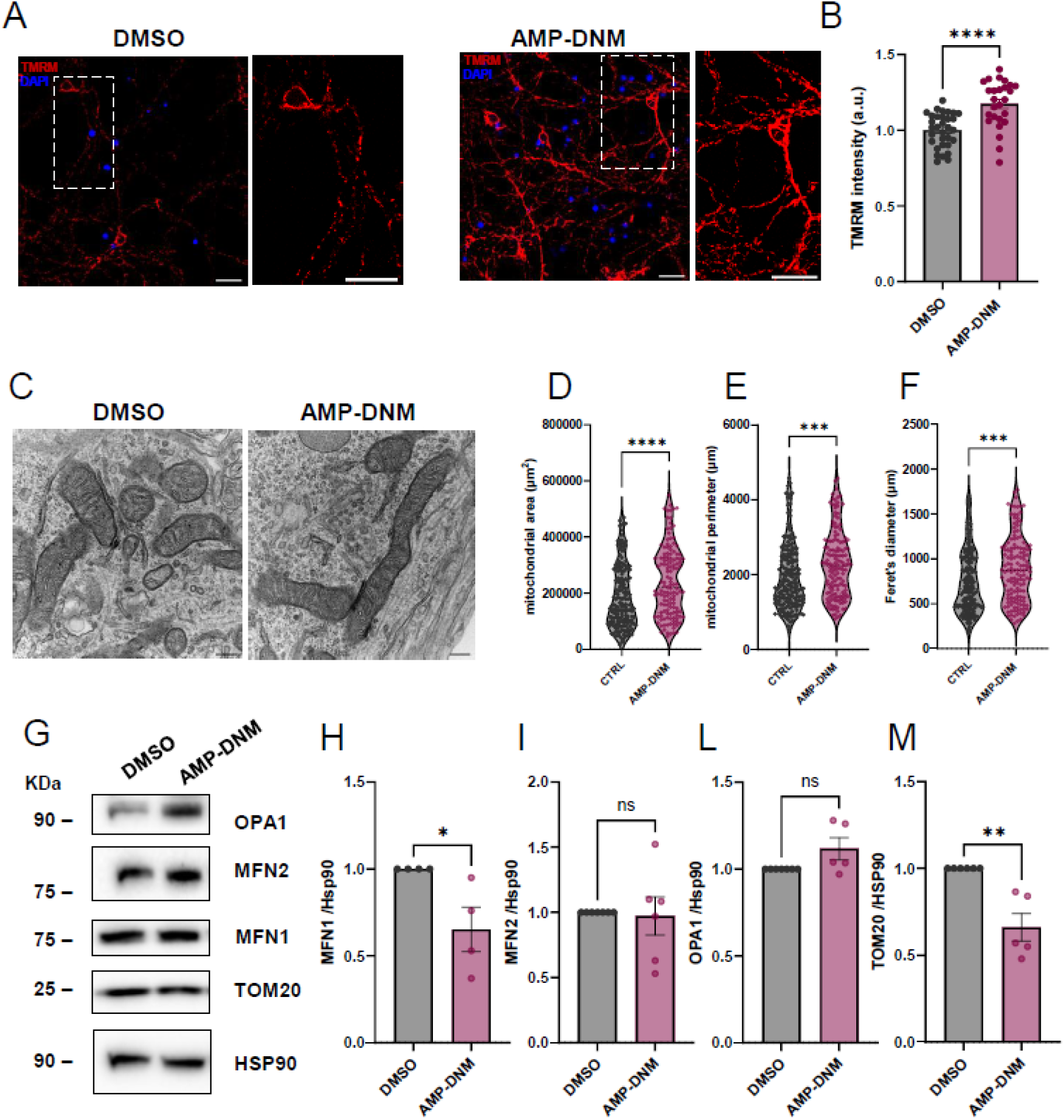
Mitochondria are altered in CGNs treated with GBA2 inhibitor. (A) Representative confocal images of TMRM staining in CGNs at 14DIV chronically treated with DMSO (0.1%) or AMP-DNM (1 nM) (scale bar=10 µm). (B) TMRM fluorescence quantification (n=30 from 4 independent experiments, Mann-Whitney *t*-test, p<0.0001). (C) Transmission electron microscopy of CGNSs at 14DIV chronically treated with DMSO (0.1%) or AMP-DNM (1nM) (scale bar=2 µm) In the magnification boxes, mitochondria with different morphologies in the two conditions (scale bar=500 nm). (D) Quantification of the mitochondrial area in GBA2-inhibition CGNs compared to controls (n=18 cell/condition from 3 independent experiments, Mann-Whitney *t*-test, p<0.0001). (E) Quantification of the mitochondrial perimeter in GBA2-inhibition CGNs compared to controls (n=18 cell/condition from 3 independent experiments, Mann-Whitney *t*-test, p=0.0003). (F) Quantification of the mitochondrial Feret’s diameter in GBA2-inhibition CGNs compared to controls (n=18 cell/condition from 3 independent experiments, Mann-Whitney *t*-test, p=0.0004). (G) Representative western blot of mitochondrial markers in CGNs at DIV14 chronically treated with DMSO or AMP-DNM. (H) MFN1 expression decreases in CGNs treated with AMP-DNM (n= 4 from 4 independent experiments, Mann-Whitney *t*-test, p=0.0286). (I) MFN2 expression is unchanged between the two conditions (n=6 from 4 independent experiments, Mann-Whitney *t*-test, p>0.9999). (L) OPA1 expression levels show no difference between AMP-DNM-treated CGNs and control (n=6 from 4 independent experiments, Mann-Whitney *t*-test, p=0.0808). (M) TOM-20 expression levels are downregulated in AMP-DNM CGNs compared to control (n=5 from 4 independent experiments, Mann-Whitney *t*-test, p=0.0022).

### GBA2 inhibition causes altered glutamate receptor function and distribution

The mitochondrial alterations in GBA2-inhibited CGNs suggested that the observed glutamate-induced Ca^2+^ dyshomeostasis is unlikely to be associated with reduced mitochondrial buffering capacity. Thus, we assessed the role of glutamate receptors in the observed Ca^2+^ signaling defects. We first performed cytosolic Ca^2+^ levels measurements using Fura-2 upon stimulation with glutamate (10 µM) in the presence or absence of NMDA or AMPA receptor antagonists (D-AP5, and NBQX, respectively) (**Fig. 4**). Baseline Ca^2+^ levels did not differ across treatment pairs (see **Additional file 1, Fig. S2**). Pharmacological inhibition of NMDA receptor (NMDAR) with D-AP5 (50 µM) increased Ca²⁺ levels in control cells (+13.8%), whereas in AMP-DNM-treated neurons it reduced cellular response, restoring near-control levels (**Fig. 4 A-C**). These findings suggest that the increase in Ca²⁺ signal observed in AMP-DNM-treated cells may depend on NMDAR activity and can be normalized upon NMDAR blockade. In contrast, antagonization of AMPA receptors with NBQX (25 µM) increased Ca²⁺ responses in both control and AMP-DNM-treated cells (+29% and +20% relative to DMSO control, respectively), but without significant differences between AMP-DNM-treated cells in the presence or absence of NBQX (**Fig. 4D-F**). These observations suggest that AMPA receptor inhibition does not normalize the AMP-DNM-associated phenotype, but rather affects Ca²⁺ responses in both conditions. The overall increased Ca^2+^ levels observed in control CGNs upon stimulation when glutamate receptors are inhibited, may be associated with the activation of inhibitory interneurons present in culture (see **Additional file 1, Fig. S3**).

**Figure 4.**
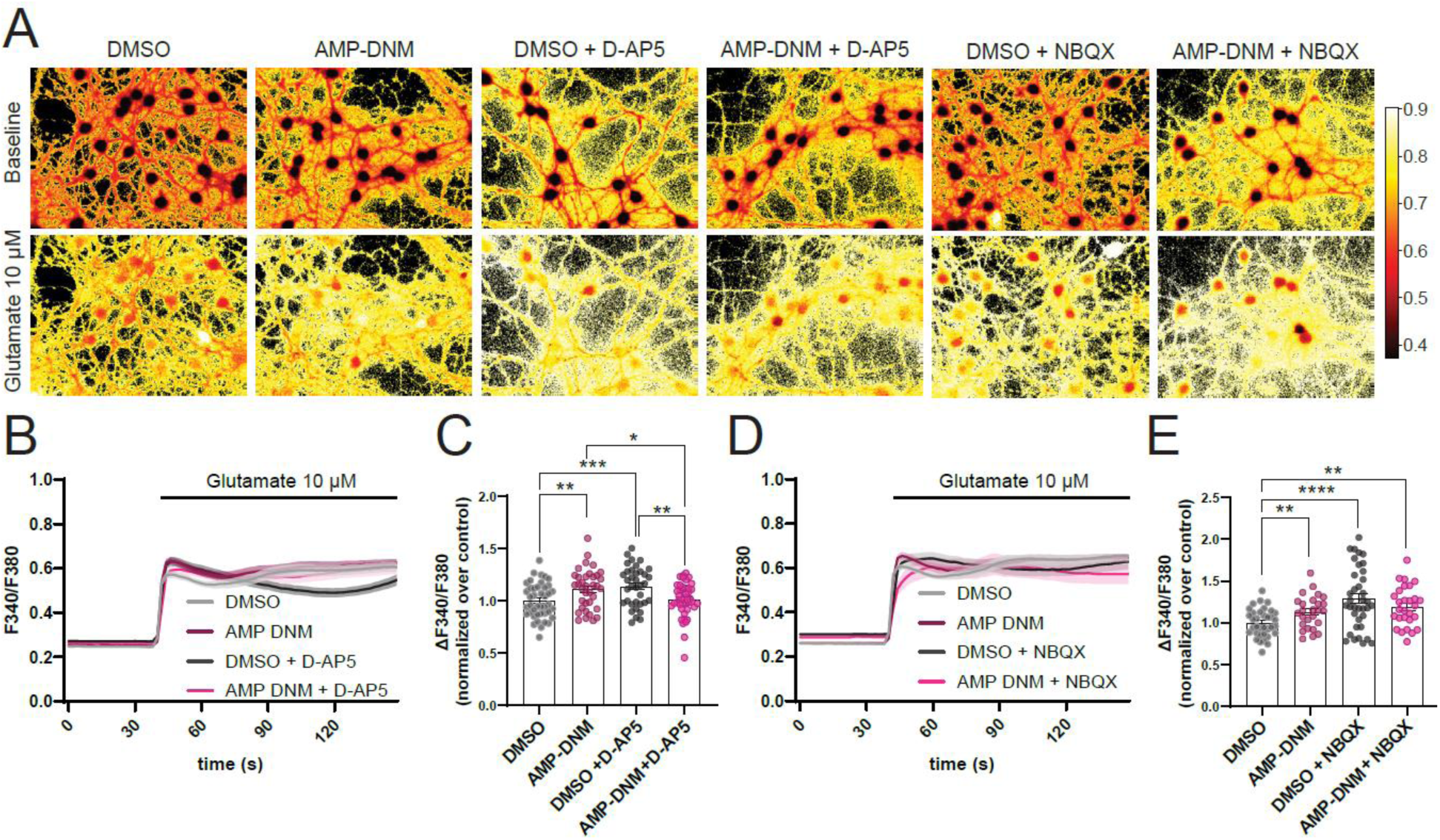
Differential contribution of NMDA and AMPA receptors to Ca²⁺ signaling in GBA2-inhibited neurons. (A) Representative ratiometric Fura-2 images (340/380 nm) of CGNs treated with DMSO or AMP-DNM (1 nM) before and after glutamate stimulation (10 µM) in the presence or absence of the NMDA receptor antagonist D-AP5 (50 µM) or AMPA receptor antagonist NBQX (25 µM). (B) Fura-2 Ca²⁺ imaging traces (± SEM) in CGNs following glutamate stimulation in the presence or absence of D-AP5 (50 µM). (C) Quantification of Ca²⁺ peak amplitude under indicated conditions (n ≥ 36, 3 independent experiments; Kruskal–Wallis test with Dunn’s multiple comparisons, uncorrected p-values; DMSO vs AMP-DNM p = 0.0078, DMSO vs DMSO + D-AP5 p = 0.0008, DMSO vs AMP-DNM + D-AP5 p=0.5767, AMP-DNM vs DMSO + D-AP5 p= 0.5086, AMP-DNM vs AMP-DNM + D-AP5 p = 0.0323, DMSO + D-AP5 vs AMP-DNM + D-AP5 p = 0.0047). (D) Fura-2 Ca²⁺ imaging traces (± SEM) in CGNs following glutamate stimulation in the presence or absence of NBQX (25 µM). (E) Quantification of Ca²⁺ peak amplitude under indicated conditions (n ≥ 26, 2 independent experiments; Kruskal-Wallis test with Dunn’s multiple comparisons, uncorrected p-values; DMSO vs AMP-DNM p = 0.0293, DMSO vs DMSO + NBQX p < 0.0001, DMSO vs AMP-DNM + NBQX p = 0.0021, AMP-DNM vs DMSO + NBQX p=0.0676, AMP-DNM vs AMP-DNM + NBQX p=0.2453, DMSO + NBQX vs AMP-DNM + NBQX p=0.5896). All data were tested for normality using the Shapiro-Wilk test.

We hypothesized that the mechanism by which NMDAR function is impaired possibly depends on an altered distribution of NMDAR at the neuronal membrane, following lipids dyshomeostasis. Thus, we first confirmed by lipidomic analysis that GlcChol levels are reduced in GBA2-inhibited CGNs (**Fig. 5A, B**), consistent with the known role of GBA2 in GlcChol synthesis [42,43]. Then, we investigated NMDAR subunit B levels and localization by western blot and confocal imaging, respectively (**Fig. 5C-F**). While total NMDAR expression was unchanged in GBA2-inhibited CGNs (**Fig. 5C, D**), its colocalization with the pre-synaptic marker VGLUT1 was increased (**Fig. 5E, F**), suggesting a functional redistribution potentially driven by GlcChol depletion. To directly test this hypothesis, CGNs were treated with exogenous GlcChol (10 μM). As we have previously demonstrated, GlcChol is taken up by the cells [13], and is expected to exert its patho-physiological functions by modulating protein activity [14].

**Figure 5.**
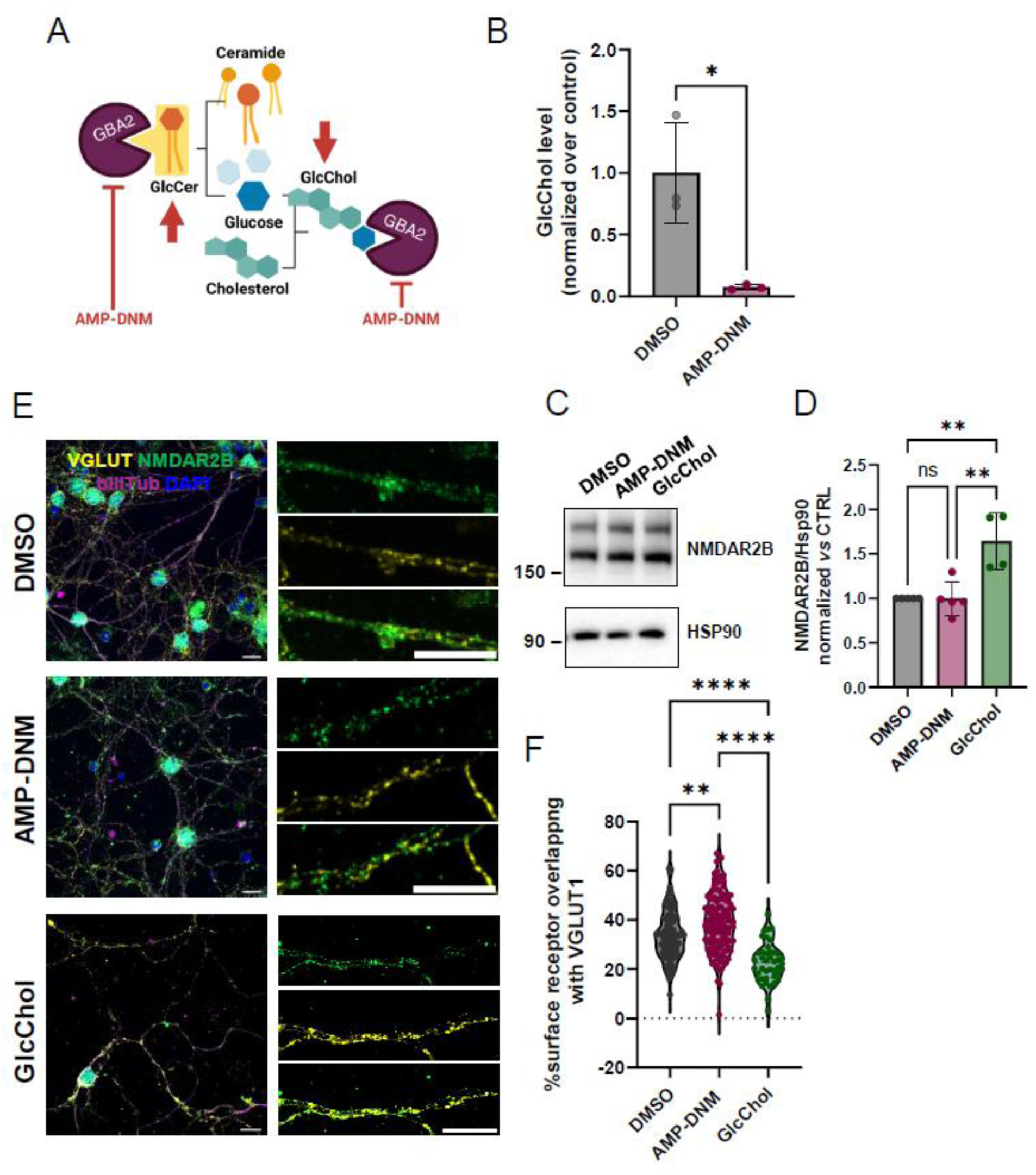
Altered lipid homeostasis following GBA2 inhibition modulates glutamate receptor localization. (A) Pharmacological inhibition of GBA2 with AMP-DNM increases glucosylceramide levels and reduces glucosylcholesterol. (B) GlcChol levels are reduced in GBA2-inhibited CGNs, as measured by LC-MS (n=3 independent experiments, unpaired *t* test; p = 0.0171). (C) Representative western blot of NMDAR2B in CGNs at DIV14 chronically treated with DMSO or AMP-DNM, or treated with GlcChol for 24 h. (D) Quantification shows that GlcChol increases NMDAR2B expression compared with both control and AMP-DNM-treated cells (n=5; mean±SEM, ordinary one-way Anova with multiple comparisons, F=14.46, p= 0.0008). (E) Representative confocal images of CGNs at DIV14 stained for VGLUT1 (presynaptic marker) and NMDAR2B (scale bar=10 µm). Arrowheads indicate sites of colocalization. (F) Quantification of colocalization between NMDAR2B and VGLUT1 in control CGNs, GBA2-inhibited CGNs and GlcChol-treated CGNs (n = 100 for DMSO, 130 for AMP-DNM, 60 for GlcChol cells from 2 independent experiments; mean±SEM; ordinary one-way Anova with multiple comparisons, F=36.78, p< 0.0001).

GlcChol treatment resulted in increased NMDAR expression by about 50% as compared to control CGNs (**Fig 5C, D**), but with reduced colocalization with VGLUT-positive structures (**Fig. 5E, F**). This is possibly due to a redistribution of NMDAR away from synaptic domains because the increased presence of GlcChol at the cell membranes could determine an alteration of lipid raft properties [44].

### GBA2 inhibition and GlcChol dyshomeostasis alter the neuronal proteome, pointing to cytoskeletal and mitochondrial dysregulation

To investigate the molecular basis of the synaptic defects observed in GBA2-inhibited CGNs, we performed proteomic profiling of GBA2-inhibited and GlcChol-treated CGNs (**Fig. 6A, B**). Altered expression of proteins involved in lipid metabolism, cytoskeletal remodeling, synaptic function and axonal growth was observed (**Fig. 6C**). In GBA2-inhibited CGNs, GBA2 protein levels were increased, consistent with the compensatory upregulation observed at earlier stages (**Fig. 1A, D**). Proteolipid protein 1 (PLP1), a structural component of myelin involved in axonal integrity maintenance [45], was reduced. EEF1AKMT4-ECE2, a readthrough protein whose C-terminal ECE2 domain has been linked to actin and microtubule cytoskeleton regulation [46], was also decreased.

**Figure 6.**
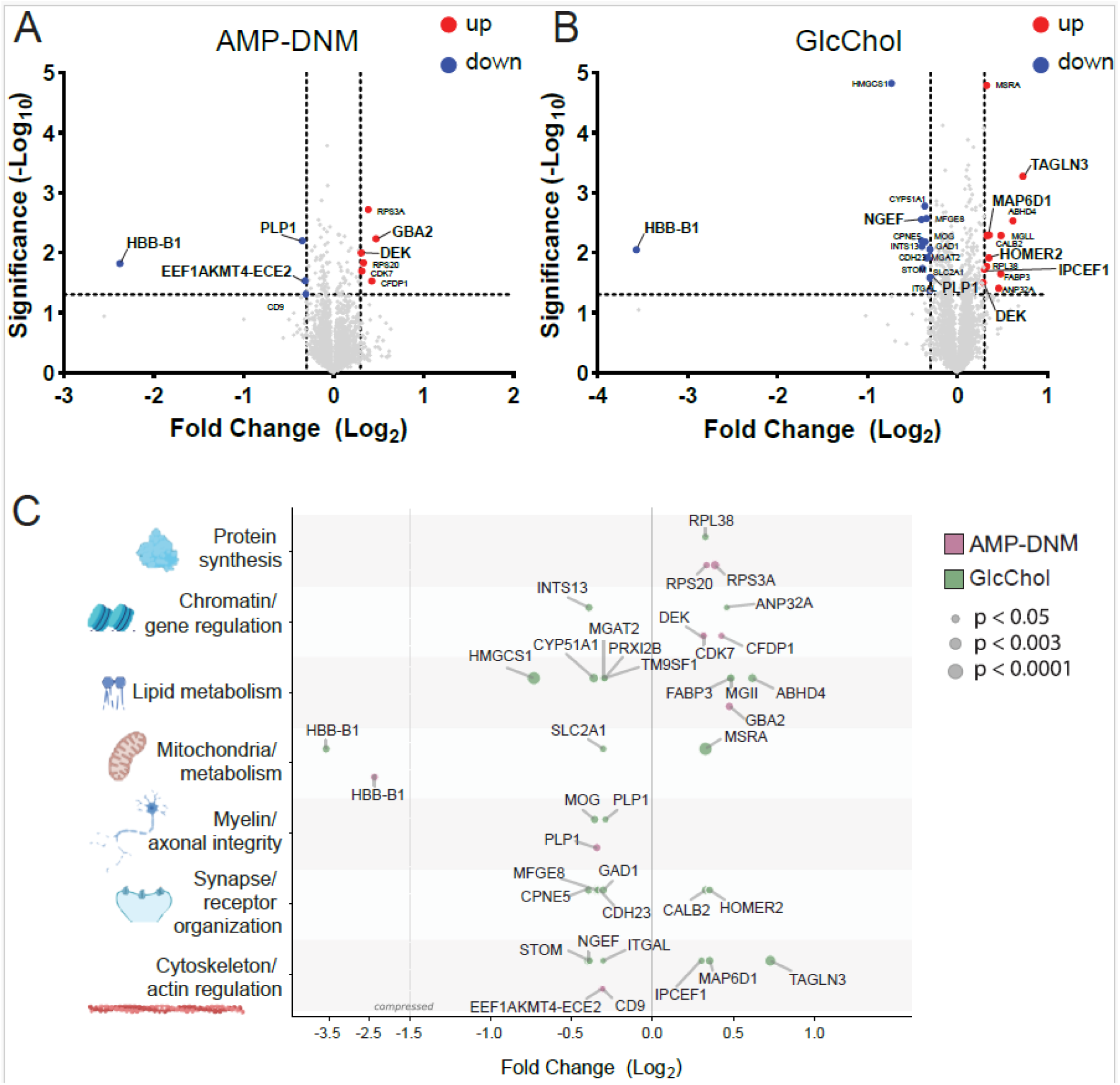
Proteome analysis shows alteration in the cytoskeletal and mitochondrial-related proteins in GBA2-inhibited neurons and GlcChol treated CGNs compared with controls. (A) Volcano plot showing differentially expressed proteins in CGNs treated with AMP-DNM (1nM) compared to control. Upregulated proteins are shown in red and downregulated proteins in blue. Dashed lines indicate significance (p-value <0.05) and log2 fold change thresholds (±0.3). Significantly altered proteins are annotated. (B) Volcano plot showing differentially expressed proteins in CGNs treated with GlcChol (10 μM) compared to control. Color coding, and thresholds are as in (A). (C) Functional classification of differentially expressed proteins grouped according to major molecular pathways, including protein synthesis, chromatin/gene regulation, lipid metabolism, mitochondrial/metabolic processes, myelin/axonal integrity, synapse/receptor organization, and cytoskeleton/actin regulation. Data are represented as log₂ fold change for AMP-DNM (purple) and GlcChol (green) treatments. Dot size reflects statistical significance (p < 0.05, p < 0.003, p < 0.0001).

DEK, a chromatin organizer involved in DNA repair and mRNA splicing, whose decrease was associated to reduced neurite development and induction of tau pathology in Alzheimer’s disease models [47,48], was instead increased by approximately 24%. Similarly, in GlcChol-treated CGNs, PLP1 was reduced by approximately 20% of control levels, and DEK was increased by about 22%, recapitulating the changes observed upon GBA2 inhibition. Additional proteins involved in actin remodeling and microtubule stabilization were altered specifically in this condition: NGEF levels were reduced, while IPCEF1, TAGLN3 and MAP6D1 were increased. HOMER2, a postsynaptic scaffolding protein, was also increased. Interestingly, the protein showing the greatest change in GBA2-inhibited CGNs and in the GlcChol-CGNs relative to controls was Haemoglobin subunit beta-1 (HBB-B1), reduced by approximately 80-90% in the two cases. HBB-B1 was shown to support mitochondrial respiration in cultured neurons [49], further pointing toward mitochondrial alterations in these models. Overall, it appears that maintaining GlcChol levels within a certain homeostatic range is key to preserving various intracellular pathways crucial for neuronal health.

### Zebrafish *gba2* crispant model recapitulates the morphological abnormalities and the motor defects of *GBA2*-HSP

To study the effects of GBA2 depletion *in vivo,* we exploited the zebrafish (ZF) model. ZF carries a single *gba2* orthologous gene on chromosome 7, which encodes a protein that shares 66% of amino acid identity and 79% of similarity with the human protein, maintaining the same enzymatic activity [50]. Analysis of *gba2* expression levels during ZF development evidenced a strong contribution from maternal mRNA, as highlighted by the high transcript concentration during the first developmental stages. Embryonic transcripts became detectable after the maternal-to-embryonic transition, increasing thereafter to reach a 7-fold expression at 5 days post fertilization (dpf) (**Fig. 7A**). Similarly to the wide expression of human *GBA2* transcripts (Protein atlas, https://www.proteinatlas.org/ENSG00000070610-GBA2/tissue), *gba2* expression was detected in all male and female adult tissues analyzed, with the higher levels in brain, gut, liver and gonads (**Fig. 7B,C**) To silence *gba2* during development, we exploited CRISPR/Cas9 technology to generate crispant ZF larvae [51], consisting of first-generation (F0) *gba2* mosaic mutants. Effective silencing was confirmed through the significant reduction of *gba2* mRNA expression (**Fig. 7D**) and of enzyme activity (**Fig. 7E**) in 5-dpf larvae. *gba1* did not compensate for this loss, since both expression levels (**Fig. 7F**) and protein activity were unaffected (**Fig. 7G**). At 5 dpf, *gba2* crispants showed increased head dimensions, measured as head length, interocular distance and eye area **(Fig. 7H-J**), bigger swim bladder area (**Fig. 7K**), but unaltered body length with respect to uninjected controls (**Fig. 7L**), suggesting possible alteration in head and brain growth. ZF larvae injected with scrambled sgRNAs did not show any morphological abnormality compared with wild type (WT) (see **Additional file 1, Fig. S4A**), nor any movement defects (see **Additional file 1, Fig. S4B-C**) thus evidencing the specificity of the observed phenotypes.

**Figure 7:**
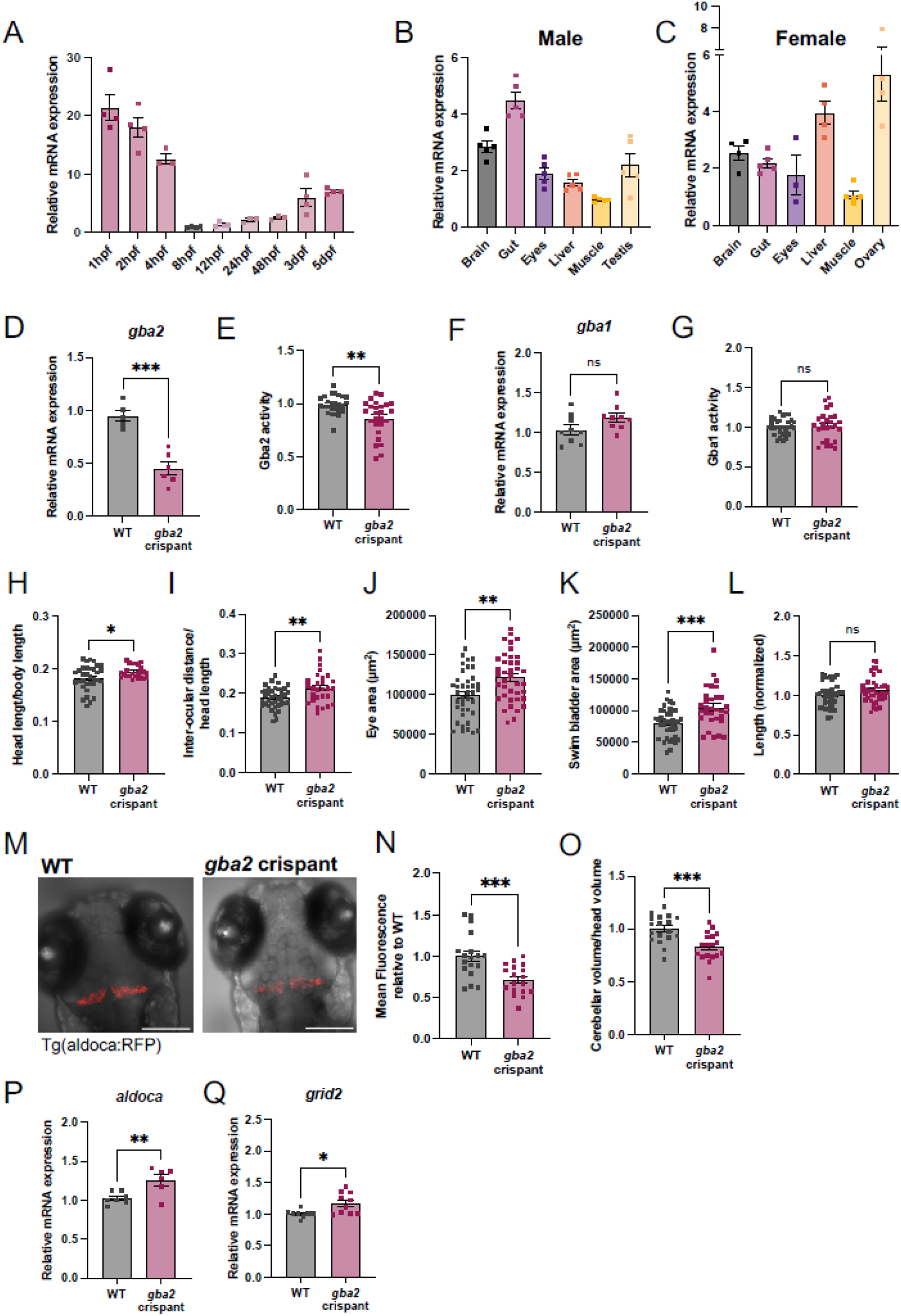
Zebrafish *gba2* crispants show reduced *gba2* expression level, head developmental alteration and cerebellar atrophy. (A) Analysis of *gba2* transcript expression in WT zebrafish embryos and larvae from 1 hpf to 5 dpf. The reference sample is 8 hpf and *ube2a* was used as the housekeeping gene. *n=*3-4 pools of embryos/larvae. (B-C) Analysis of *gba2* transcript abundance in 4-6 mpf zebrafish male (B) and female (C) tissues. Muscle was considered as the reference sample, and *β-actin* was used as the housekeeping gene. *n=*3-5 tissue samples from WT animals. (D) Analysis of *gba2* transcript expression (*n≥*5 pools of 5-dpf zebrafish larvae from 4 independent experiments, unpaired Student’s *t-*test, p=0.0002). (E) Analysis of Gba2 protein activity in WT and *gba2* crispants (*n*=26 pools of 5-dpf zebrafish larvae from 4 independent experiments, Welch’s *t-*test, p=0.0011). (F) Analysis of *gba1* transcript expression (*n*=9 pools of 5-dpf zebrafish larvae from 4 independent experiments, unpaired Student’s *t-*test, p=0.0766). (G) Analysis of Gba1 protein activity in WT and *gba2* crispants (*n≥*26 pools of 5-dpf zebrafish larvae from 4 independent experiments, Welch’s *t-*test, p=0.7233). (H-L) Morphometric analysis of 5-dpf WT and *gba2* crispant larvae (*n≥*29 from 3 independent experiments, unpaired Student’s *t-*test for standard length, p=0.0661 (H), swim bladder area, p=0.0003 (I) and eye area, p=0.0019 (J); Welch’s *t-*test for head length/body length, p=0.0188 (K) and interocular distance/head length, p=0.0051 (L)). (M) Representative confocal images of transgenic Tg(aldoca:RFP) 5-dpf WT and *gba2* crispant zebrafish larvae (scale bar=200µm). (N) Quantification of the mean fluorescence in RFP-*aldoca*-positive Purkinje cells in 5-dpf *gba2* crispants relative to WT (*n*=18 from 3 independent experiments, unpaired Student’s *t-*test, p=0.0008). (O) Measure of cerebellar volume normalized for the head volume (*n*=18 from 3 independent experiments, unpaired Student’s *t-*test, p=0.0005). (P-Q) Relative mRNA expression of cerebellum-related genes (*n≥*6 pools of 5-dpf zebrafish larvae from 4 independent experiments, unpaired Student’s *t-*test for *aldoca*, p=0.009 (P); Welch’s *t-*test for *grid2*, p=0.0107 (Q)).

Given that patients with SPG46 often present cerebellar atrophy [4,52], we exploited the Tg(aldoca:epNTR-TagRFP)^nub125^ transgenic line (hereafter Tg(aldoca:RFP)) [26], which specifically labels Purkinje cell bodies, to verify any difference in cerebellar development and size (**Fig. 7M**). At 5 dpf, we observed a significant reduction in the mean RFP fluorescence in *gba2* transgenic crispants compared to controls (**Fig. 7N**) and an associated reduction of cerebellar volume (**Fig. 7O**). Intriguingly, aldolase C, fructose-bisphosphate (*aldoca*) gene expression was higher in *gba2* crispants (**Fig. 7P**). In line with this, we also observed increased expression of glutamate receptor d2 (*grid2*) mRNA (**Fig. 7Q**), that encodes a protein specifically expressed in the dendritic spines of Purkinje cells. Grid2 regulates synaptic organization and plasticity, thus its potential overexpression, together with increased *aldoca,* could indicate higher spine density at the dendrites, as a compensatory response of stressed Purkinje neurons.

Coherently with movement defects reported in HSP patients, we observed that *gba2* crispants show impaired swimming performances when exposed to alternate light-dark stimuli, with a significant reduction in the distance moved during both light and dark phases (**Fig. 8A, B**). Analysis of trunk skeletal muscles, however, only evidenced a mild reduction in the birefringence index, without structural breakdown of muscle fibers (**Fig. 8C, D**), supporting the hypothesis that movement alteration may be mainly ascribed to neurological damage more than muscular weakness.

**Figure 8:**
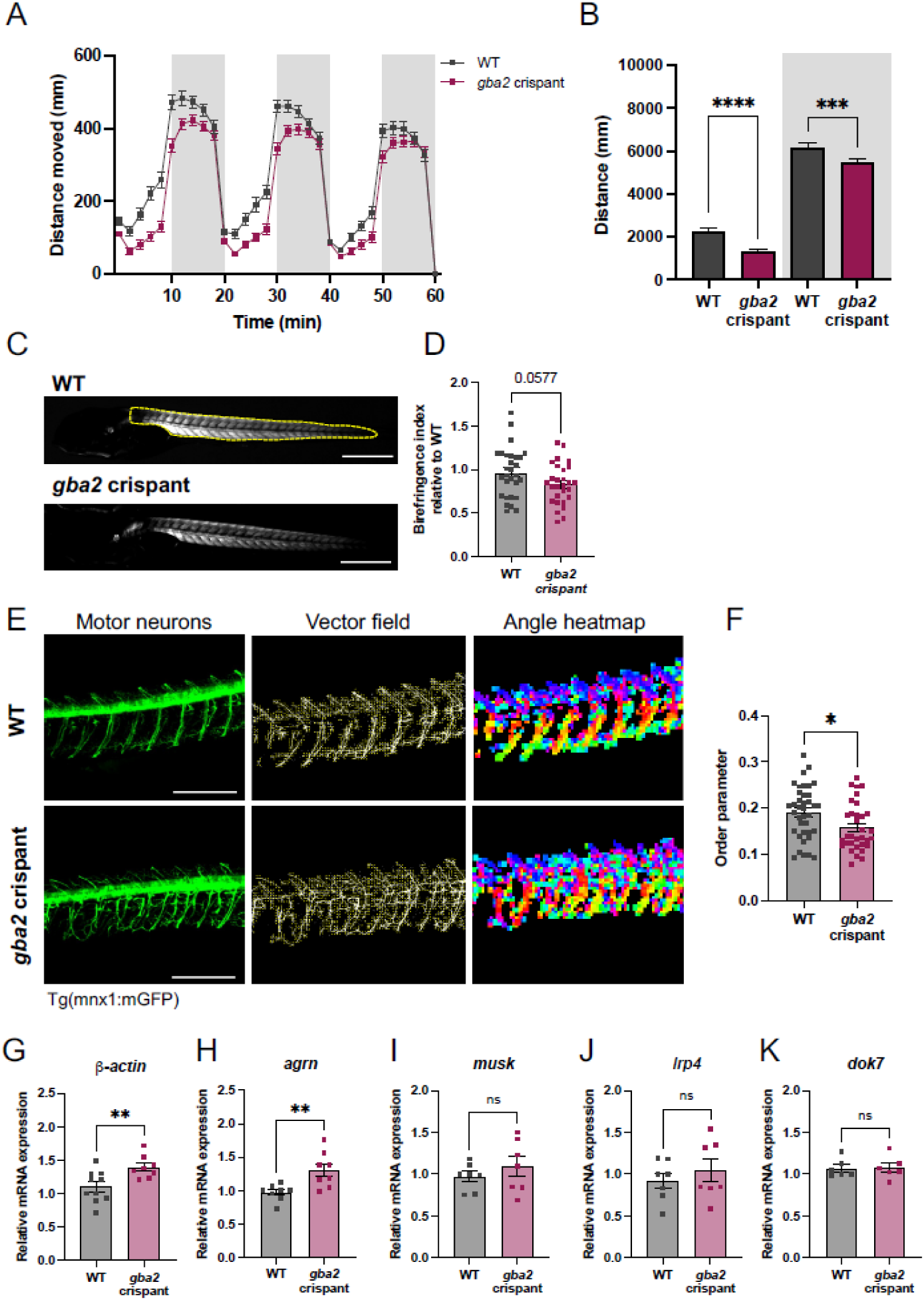
Zebrafish *gba2* crispants show impaired locomotor behavior and motor neuron defects. (A) Analysis of swimming behavior after alternating light (white background)/dark (grey background) cycles stimulation. Data are presented as the mean distance covered over time by *n=*120 5-dpf zebrafish larvae from 5 independent experiments. (B) Quantification of the distance moved during the light (white background) and dark (grey background) periods (*n=*120 from 5 independent experiments, Mann-Whitney test, p<0.0001 during light phase, p=0.0006 during dark phase). (C) Representative photographs of 5-dpf zebrafish larvae under polarized lenses (scale bar=500µm). Analysis of skeletal muscle birefringence index was performed in the outlined trunk region. (D) Analysis of birefringence index (*n*=30 from 3 independent experiments, unpaired Student’s *t-*test, p=0.0577). (E) Representative confocal images of transgenic Tg(mnx1:mGFP) 3-dpf WT and *gba2* crispants (left panel, scale bar=200µm) flanked by vector field visualization (middle panel) and angle heatmap representation (right panel). F) Quantification of the “Order parameter” (*n*≥30 from 3 independent experiments, Mann-Whitney test, p=0.0161). (G-K) Relative mRNA expression of cytoskeleton- and NMJ-related genes (*n≥*6 pools of 5-dpf zebrafish larvae from 4 independent experiments, unpaired Student’s *t-*test for *actin*, p=0.0094 (G), *agrn,* p=0.0057 (H), *musk,* p=0.3877 (I), *lrp4,* p= 0.4567 (J), *dok7,* p=0.9639 (K)).

### Cytoskeletal abnormalities in motor neurons and neuromuscular junctions are also observed in zebrafish *gba2* crispants

Considering the defects observed in mice CGNs and the motor impairment observed in ZF crispants, possibly associated with neuronal dysfunction, we further investigated this aspect *in vivo*.

While ZF lack true corticospinal neurons, which are mainly affected in HSPs [53], spinal motor neurons provide a well-established *in vivo* model to study the stability of long-projecting axons. These neurons share key structural features with corticospinal neurons, including extreme axonal polarization and dependence on cytoskeletal and membrane dynamics. Thanks to the transgenic zebrafish line Tg(mnx1:mGFP), which specifically labels motor neurons, we measured axonal branching in *gba2* crispants as compared to control ZF. To quantify the alignment of different fragments of motor neuron growth, we employed a Fourier-Transformed (FT)-based approach [36] that yields a numerical “order parameter”, describing the degree of complexity of the neuronal network (**Fig. 8E**). At 3 dpf, when the axonal complexity is still reduced, *gba2* crispants show a significantly lower “order parameter” compared to controls, consistent with a disorganized and chaotic terminal arborization (**Fig. 8F**). We linked this scenario with an abnormal branching, in line with previous literature [9,17]. Consistently, by RT-qPCR we observed higher expression of *β-actin*, a main cytoskeletal component (**Fig. 8G**), together with *agrin* (*agrn*) (**Fig. 8H**), which encodes a protein secreted by motor neurons to recruit acetylcholine receptors (AchR) to the neuromuscular junction (NMJ) and form functional synapses [54]. Expression of *muscle associated receptor tyrosine kinase* (*musk*)*, low-density lipoprotein receptor-related protein 4* (*lrp4*) and *docking protein 7* (*dok7*), the main components of the NMJ post-synaptic compartment, were not affected by *gba2* silencing (**Fig. 8I-K**), pointing toward the hypothesis that defective cytoskeletal organization due to alterations in the neuronal membranes may impact receptors distribution at the synapse, rather than impairing the expression of key proteins for the post-synaptic compartment.

### Mitochondria alterations are also present in ZF *gba2* crispants

To verify the presence of mitochondrial defects in this model, we evaluated the expression of mitochondrial-related genes in *gba2* crispants at 5dpf. We found reduced *fission, mitochondrial 1* (*fis1*) expression levels (**Fig. 9A**) and increased *mitofusin 2* (*mfn2*) in *gba2* crispants compared to WT (**Fig. 9B**). This points to a scenario of altered mitochondrial remodeling, consistent with the results obtained by TEM in cultured mouse cerebellar neurons. The fact that *opa1* transcripts remain unchanged (**Fig. 9C**), as well as other genes involved in mitochondrial respiration (*NADH:ubiquinone oxidoreductase complex assembly factor 1*, *ndufaf1*) (**Fig. 9D**) and ROS management (mitochondrial *superoxide dismutase 2, sod2* and cytoplasmic *sod1*) (**Fig. 9E,F**), suggests that, at least at this time-point, larvae do not experience defective mitochondrial respiration and oxidative stress. The significant increase of *hypoxia-inducible factor 1a-3a* (*hif1a-3a*) (**Fig. 9G**) and *erythropoietin* (*epo*) (**Fig. 9H**), both involved in the regulation of cellular response to hypoxia, is consistent with the reduction of HBB-B1 subunit levels observed by proteomics in mouse CGNs, which supports the hypothesis of initial defects in mitochondrial function. Moreover, the increase of TMRM signal in *gba2* crispants measured by confocal microscopy in ZF heads (**Fig. 9I-J**), suggests altered mitochondrial functionality or could be due to the presence of enlarged mitochondria.

**Figure 9:**
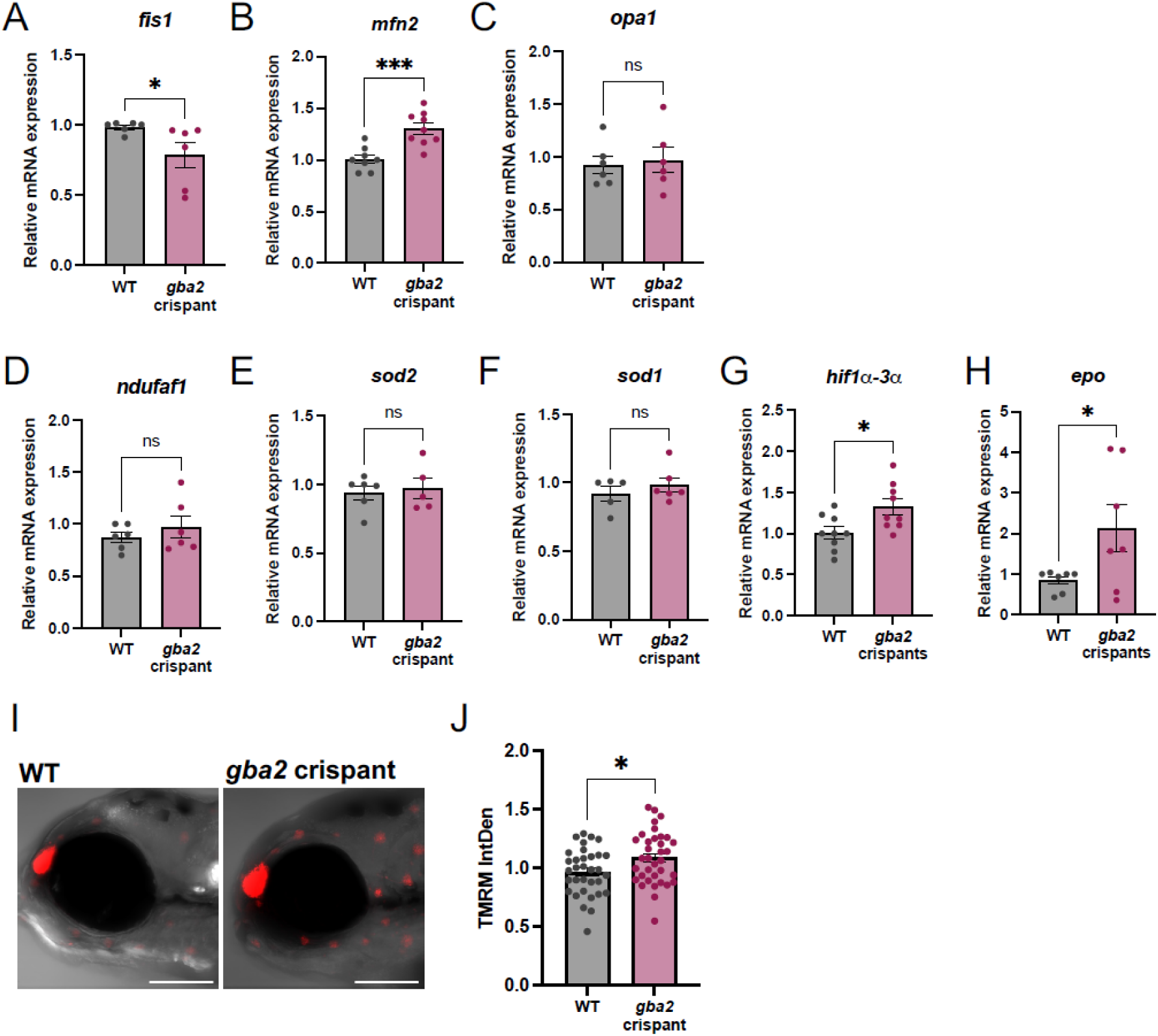
Zebrafish *gba2* crispants present altered mitochondria. (A-H) Relative mRNA expression of mitochondria- and hypoxia-related genes (*n≥*6 pools of 5-dpf zebrafish larvae from 4 independent experiments, Mann-Whitney for *fis1*, p=0.0108 (A), unpaired Student’s *t-*test for *mfn2,* p=0.0006 (B), *opa1,* p=0.7636 (C), *ndufaf1,* p=0.4142 (D), *sod2,* p=0.7383 (E), *sod1,* p=0.4361 (F), *hif1α-3α,* p=0.0171 (G), *epo,* p=0.0487 (H)). (I) Representative confocal micrographs of 5-dpf WT and *gba2* crispant zebrafish larvae stained with TMRM (scale bar=200µm). (J) Quantification of TMRM integrated density relative to WT (*n≥*33 from 3 independent experiments, unpaired Student’s *t-*test, p=0.0232).

### GBA2-HSP patient-derived fibroblasts recapitulate the mitochondrial dysfunction

To support translational significance of the obtained results, we investigated mitochondrial function, cytoskeletal and RAC1 impairment in human-derived cells from an HSP-*GBA2* patient. Skin fibroblasts were derived from a patient carrying the homozygous c.2063G>A (p.Cys688Thr) *GBA2* mutation presenting an early onset HSP form, characterized by spasticity, cerebellar syndrome, mild cognitive impairment and cataract [4]. Consistent with most HSP-associated *GBA2* variants [9], this mutation is located within the catalytic domain of the enzyme.

We first assessed GBA2 protein levels by immunofluorescence analysis (**Fig. 10A, B**), which showed a reduction of about 50%, likely associated with increased degradation of the mutated form. Enzymatic activity assays performed on fibroblast lysates revealed a 40% reduction in GBA2 activity (**Fig. 10C**), in line with the observed reduction in the protein amount, but also possibly associated with the reduced enzymatic activity due to the mutation occurring in GBA2 catalytic domain.

**Figure 10:**
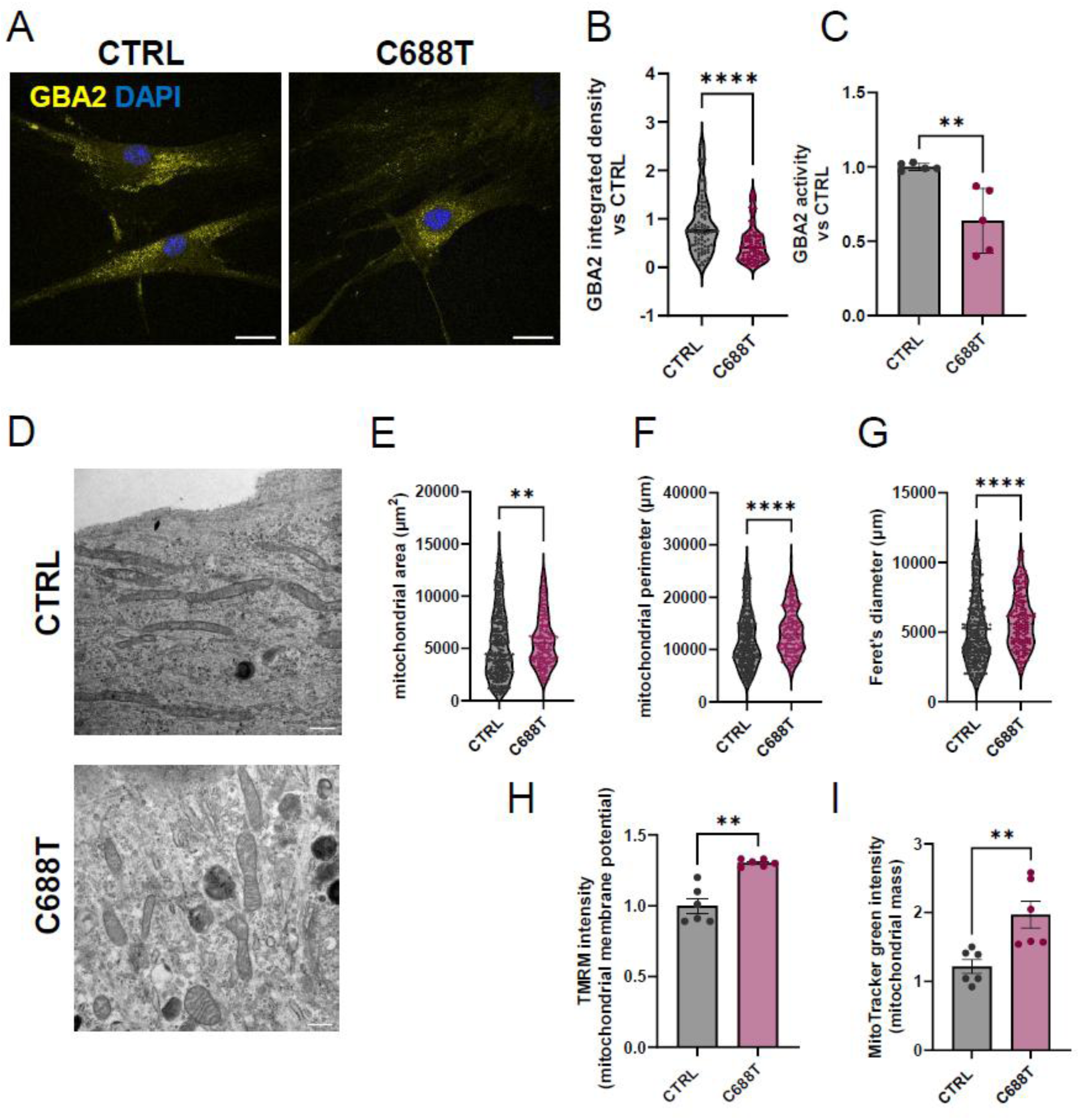
C688T mutation decreases GBA2 expression and activity and alters mitochondrial morphology in human fibroblasts. (A) Representative confocal images of human fibroblasts from healthy (CTRL) and HSP patients carrying the c.2063G>A (p.Cys688Thr) mutation (C688T) stained with GBA2 antibody (scale bar=30 um). (B) C688T fibroblasts reported significantly lower GBA2 integrated density compared to CTRL (n= 14 cells (7 CTRL and 7 C688T) from 2 independent experiment, mean± SEM, Mann-Whitney *t*-test, p<0.0001). (C) GBA2 activity decreased in mutated fibroblast compared to control (n=6 from 2 independent experiments, mean± SEM, Mann-Whitney *t*-test, p=0.079). (D) Transmission electron microscopy of human fibroblasts from healthy patients (CTRL) and carrying GBA2 mutation (C688T) (scale bar=2 µm). GBA2 mutation increases mitochondrial area (E), perimeter (F) and Feret’s diameter (G) (n=5cells/condition, mean± SEM, Mann-Whitney *t*-test, p=0.096 for Area, p<0.0001 for Feret’s diameter, p<0.0001 for Perimeter). (H) Mitochondrial membrane potential increased in C688T fibroblasts evaluated by TMRM assay by flow-cytometry (n=6 from 2 independent experiments, Mann-Whitney *t*-test, p=0.0022); (I) Mitotracker green fluorescence, evaluated by flow-cytometry, significative increase in C688T fibroblast confirming increase in mitochondrial mass (n=6 from 2 independent experiments, Mann-Whitney *t*-test, p=0.0022).

Similarly to what was observed in murine GBA2-inhibited CGNs, TEM analysis of GBA2^Cys688Thr^ mutant fibroblasts (**Fig. 10D**) revealed significant increase in mitochondrial area, perimeter, and Feret’s diameter compared to controls (**Fig. 10E-G**). Interestingly, mitochondrial membrane potential was altered also in mutant GBA2^Cys688Thr^ fibroblasts, as indicated by increased TMRM fluorescence measured by flow cytometry (**Fig. 10H**). In addition, MitoTracker fluorescence, used as a marker of mitochondrial mass, was also increased (**Fig. 10I**). Overall, these results are coherent with the data obtained in the neuronal and zebrafish models, where the mitochondrial compartment is enlarged, probably undergoing altered fusion-fission dynamics, thus establishing their relevance for the human disease context.

### RAC1 is modulated by GlcChol in GBA2-linked models

Previous reports showed that the activity of RAC1 was increased in GBA2-KO murine fibroblasts, and associated with cytoskeletal impairment and disorganization of neuronal structures [18], as we recapitulated in our models. Mitochondrial fusion/fission balance is also known to be regulated by cycling of actin filaments onto and off of mitochondria, requiring dynamic actin polymerization controlled by RAC1 downstream targets [55] (**Fig. 11A**). Therefore, we directly assessed whether RAC1 level and function were altered in our GBA2-HSP models, and if this could be associated with GlcChol dyshomeostasis. We first analyzed the expression of proteins involved in RAC1-dependent actin cytoskeleton regulation in GBA2-inhibited and GlcChol-treated CGNs at DIV 14 (**Fig. 11B**). The phosphorylation of RAC1 direct substrate, PAK1, only showed trends towards an increase in the GBA2-inhibited CGNs, and towards a decrease in the GlcChol-treated cells, although not significant (**Fig. 11C**). Phospho-COFILIN, which is the inactive form of the protein and a direct substrate of PAK1, was significantly reduced in GlcChol-treated CGNs (**Fig. 11D**), suggesting at least a mild reduction in RAC1 activity. For the GBA2-inhibited CGNs, these results may be associated with the compensatory mechanisms we already observed at the morphological and protein levels. In the neuronal GBA2-HSP model, we observed a significantly increased (+20%) RAC1 expression level, whereas the GlcChol treatment caused a reduction of RAC1 protein of about 20% (**Fig. 11E**). Interestingly, the levels of the post-synaptic structural actin-binding protein DREBRIN [56] were unchanged in both conditions (**Fig. 11F**), suggesting that the structural integrity of the post-synaptic actin scaffold is largely preserved. Thus, the alterations in HOMER2 levels and NMDAR distribution we observed, may reflect a functional reorganization of post-synaptic receptor complexes rather than a collapse of post-synaptic architecture. Coherently with the proteomics data (**Fig. 6**), the levels of TAGLN, a neuronal protein fundamental in the development of the central nervous system that acts stabilizing the actin cytoskeleton [57], were significantly increased in GlcChol-treated CGNs (**Fig. 11G**), further suggesting that GlcChol can modulate this process.

**Figure 11:**
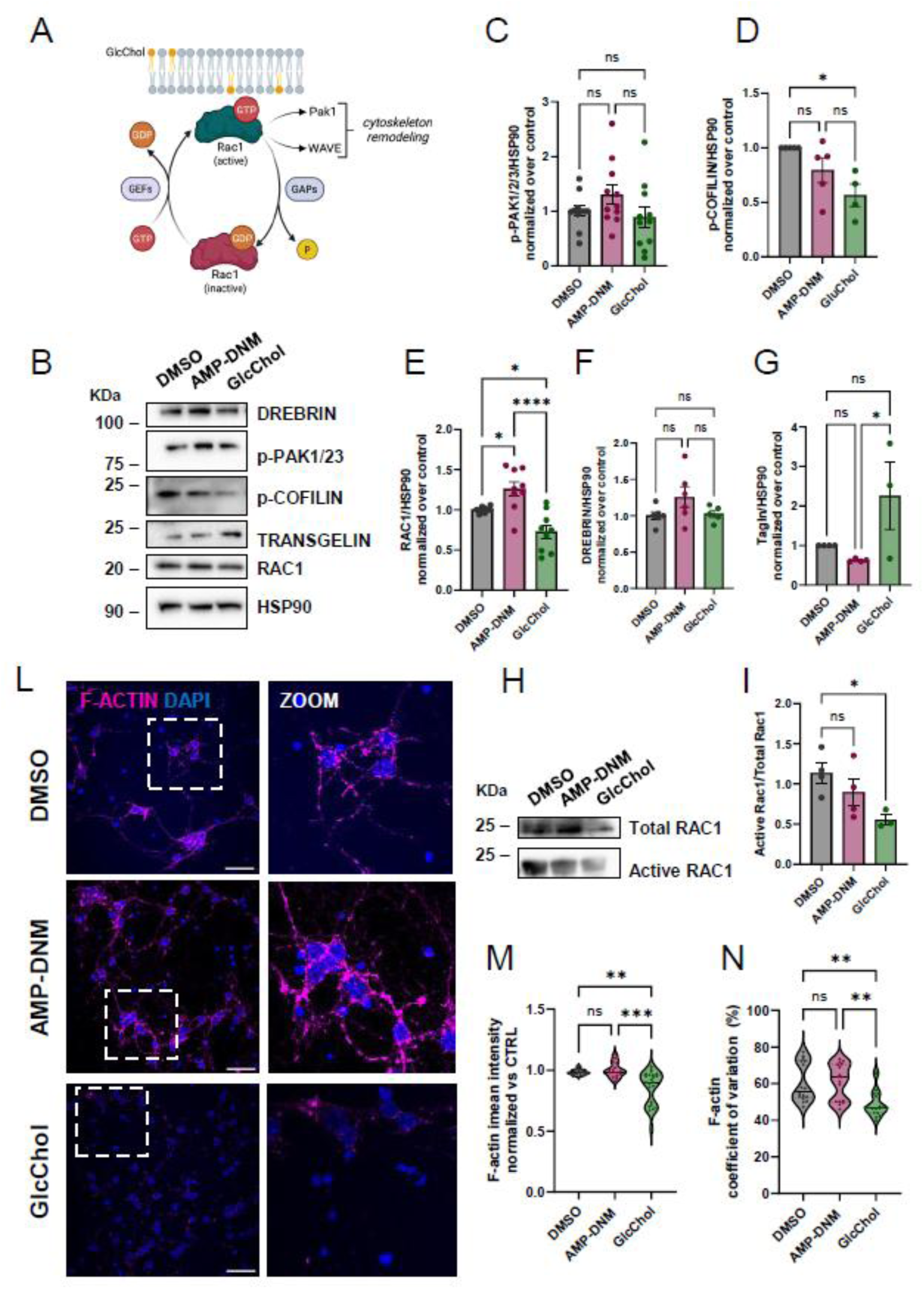
Rac1 activity and distribution impact on GBA2-inhibited CGNs. (A) Rac1 acts as a molecular switch, alternating between an inactive GDP-bound form (primarily in the cytoplasm) and an active GTP-bound form that is located at the plasma membrane. When activated, Rac1 translocates to the plasma membrane to regulate cytoskeleton remodeling, by modulating its effectors, such as PAK1 and WAVE. GEFs (Guanine Nucleotide Exchange Factors) and GAPs (GTPase-activating proteins) are involved in the switching between the active and inactive state of Rac1. (B) Representative western blot of DREBRIN, p-PAK1/2/3, p-COFILIN, TAGLN and RAC1 in CGNs at DIV14 chronically treated with DMSO or AMP-DNM, or treated with GlcChol for 24 h. (C) No significant changes were observed on the expression of p-PAK1/2/3 direct substrate of RAC1 (n=11 from 7 independent experiment, mean±SEM, ordinary one-way Anova with multiple comparisons, F=1.839, p=0.1764. (D) p-COFILIN expression decreases in GlcChol-treated CGNs compared with CTRL (n=5 from 4 independent experiment, mean±SEM, ordinary one-way Anova with multiple comparisons, F=5.910, p=0.0181). (E) Quantification shows that RAC1 increases in AMP-DNM inhibited CGNs and decreases in GlcChol treated CGNs compared with CTRL (n=10 from 7 independent experiment; mean±SEM, ordinary one-way Anova with multiple comparisons, F=14.47, p< 0.0001); (F) No changes were observed in the expresison of DREBRIN (n=6 from 4 independent experiment, mean±SEM, ordinary one-way Anova with multiple comparisons, F=2.238, p=0.1434). (G) Transgelin (TAGLN) expression significative increases in GlcChol treated CGNs compared with both CTRL and AMP-DNM (n=4 from 4 independent experiment, mean±SEM, ordinary one-way Anova with multiple comparisons, F=4.440, p=0.0504); (H) Representative western blot of total RAC1 and active-RAC1 isolated from neuronal lysated by pull-down assay; (I) Quantification of RAC1 activity reported as the ratio of active-RAC1 to total and normalized to control conditions (n=4 from 3 independent experiments, mean±SEM, ordinary one-way Anova with multiple comparisons, F=4.050, p=0.0610). (L) Representative confocal images of CGNs at DIV14 stained for F-ACTIN and β-III-Tubulin (scale bar=10 µm). (M) F-ACTIN mean intensity as well as coefficient of variation (N) decreases in GlcChol treated CGNs compared with both CTRL and AMP-DNM (n= 8/condition from 2 independent experiment, mean±SEM, ordinary one-way Anova with multiple comparisons Kruskal-Wallis test, p<0.0001 for mean intensity and p= 0.0004 for coefficient of variation).

To better understand the role of RAC1 in this model, we evaluated RAC1 activity by a pull-down assay in the CGNs model. Active RAC1 (RAC1-GTP) was isolated (see **Additional file 1, Fig. S5**; **Fig. 11H**), but no changes were observed in AMP-DNM treated CGNs as compared to controls, whereas GlcChol causes a significant reduction in active RAC1 levels (**Fig 11I**).

To further explore the role of RAC1 in GBA2-inhibited and GlcChol-treated CGNs, we evaluated F-ACTIN levels and distribution within the cells (**Fig. 11L**), to quantify the downstream effects of the RAC1 modulation on actin cytoskeleton polymerization and organization. GlcChol treatment decreased F-ACTIN mean fluorescence intensity (**Fig. 11M**) compared to both control and AMP-DNM-treated CGNs. Moreover, F-ACTIN coefficient of variation was decreased in GlcChol-treated CGNs (**Fig. 11N**), suggesting an overall loss and more homogeneous distribution of polymerized actin in this condition. Together, these findings suggest an impairment of RAC1-dependent actin regulatory signaling upon acute GlcChol treatment, altering actin cytoskeletal remodeling. Conversely, global RAC1 activity seemed not altered upon GBA2 chronic inhibition, which may suggest that it may be altered only at a specific cell compartment, i.e. the synapse, in which we observe neuronal defects associated to cytoskeletal disorganization.

To provide translational support to this finding, we studied RAC1 and F-ACTIN cytoskeleton in the patient-derived mutant GBA2^Cys688Thr^ fibroblast line. Interestingly, RAC1 fluorescence intensity and the coefficient of variation of the signal (**Fig. 12A-C**), which measure the distribution of the protein within the cell, were decreased in GBA2^Cys688Thr^ patient fibroblasts. These data point toward a more diffuse spreading of the protein within the cell rather than a reduction in RAC1 expression. To further investigate the downstream effect of RAC1 alteration on actin filaments in this model, we evaluated F-ACTIN signal intensity and organization by confocal imaging (**Fig. 12A**). While the overall F-ACTIN fluorescence intensity and the coefficient of variation of the intracellular F-ACTIN fluorescence signal were comparable between patients and controls (**Fig. 12D-E**), indicating similar overall protein levels and distribution, patients showed a significantly higher skewness (**Fig. 12F**), suggesting an asymmetric redistribution of the signal toward focal areas of increased intensity within individual cells, consistent with altered actin polymerization dynamics.

**Figure 12:**
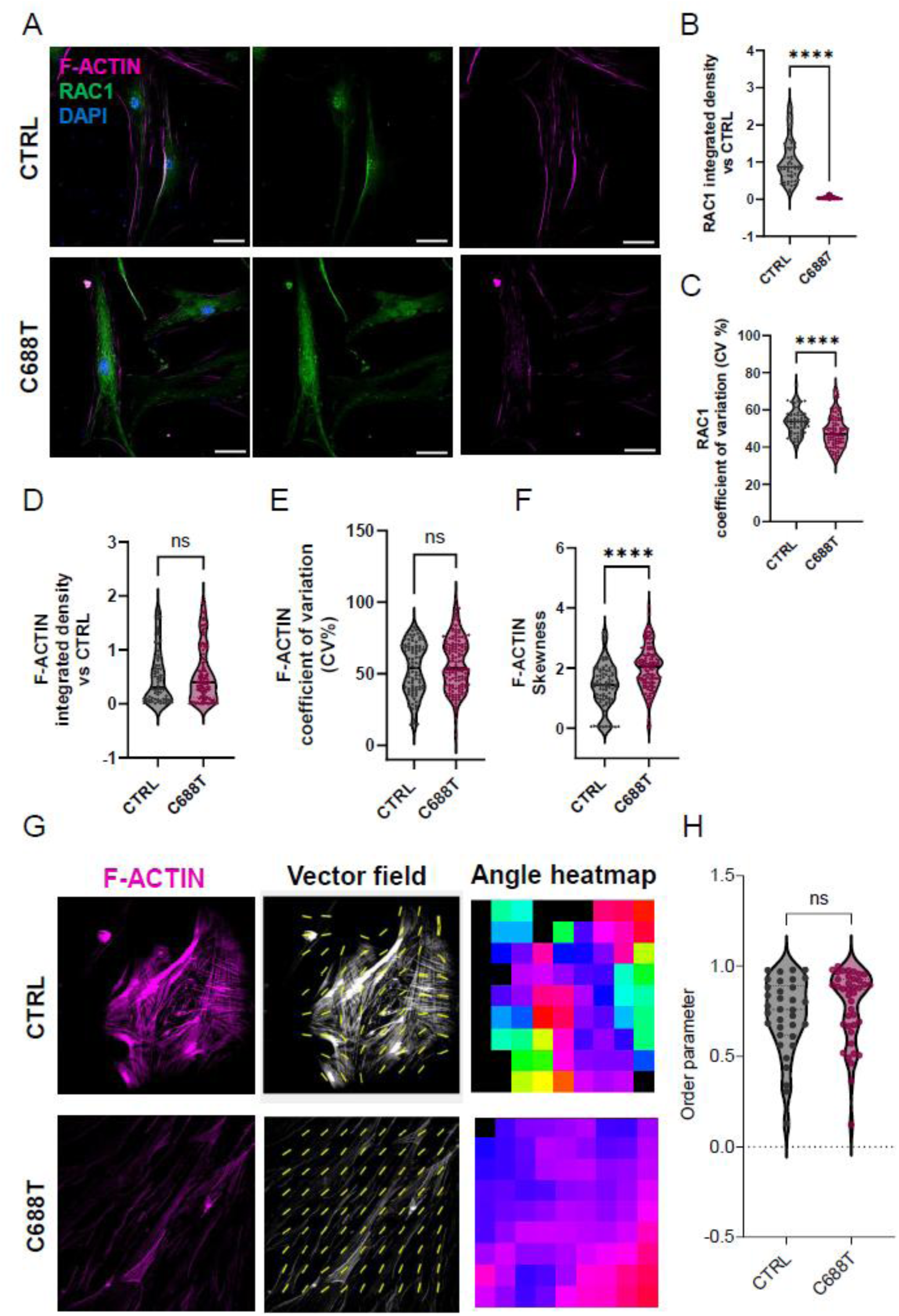
C688T mutant fibroblast recapitulates cytoskeletal disorganization. (A) Representative confocal images of healthy and C688T fibroblasts stained for F-actin and RAC1 (scale bar=10µm); (B) RAC1 integrated density is significantly reduced in C688T fibroblasts compared to control along with coefficient of variation (n= 3 independent experiment, mean±SEM, Mann-Whitney t-test, p<0.0001)(C); (D) F-ACTIN integrated density and coefficient of variation (E) show a non-significant increasing trend in C688T fibroblasts, and (F) F-ACTIN fluorescence signal skewness is increased in C688T fibroblasts as compared to controls (n= 3 independent experiment, mean±SEM, Mann-Whitney t-test, p=0.5784 for integrated density, p=0.1622 for coefficient variation, p<0.0001 for skewness (G) Representative actin alignment analysis performed using the Alignment by Fourier Transform (AFT) algorithm, Local fiber orientation was determined by FFT-based analysis of overlapping image windows, generating orientation vector fields (yellow) and orientation heatmaps displaying the spatial distribution of fiber orientation. (H) Violin plot showing the median local orientation order parameter for CTRL and C688T fibroblasts. No significant differences were detected (n= 3 independent experiment, mean±SEM, Mann-Whitney t-test, p=0.2197).

To further evaluate the organization of actin fibers within control and patient fibroblast, we exploited the FT algorithm previously used to measure ZF motor neurons organization (**Fig. 8E**) and already applied to this type of signals [36]. The FT analysis suggested that F-ACTIN filaments in GBA2-HSP patient cells are more ordered, which may be coherent with a reduced amount of branched actin due to RAC1 altered distribution.

### RAC1 inhibition rescues motor and molecular phenotypes in ZF *gba2* crispants

To verify whether the reduction of RAC1 activity could rescue the pathological phenotypes observed in *in vivo* model, we treated *gba2* crispants and WT zebrafish larvae with RAC1 inhibitor NSC23766, at a final concentration of 10 µM [58]. After 24 hours of treatment, we documented a complete recovery of axonal network organization (**Fig. 13A**), together with the restoration of near-control expression levels of motor neuron-secreted *agrn* (**Fig. 13B**), of cerebellar *aldoca* (**Fig. 13D**) and *grid2* (**Fig. 13E**), and an indication toward the recovery of mitochondrial dynamics concerning *mfn2* (**Fig. 13C**). Moreover, we found a significant improvement of motor activity (**Fig. 13F-G**), indicating RAC1 as a promising target to be further investigated as a possible therapeutic strategy in *GBA2*-HSP.

**Figure 13:**
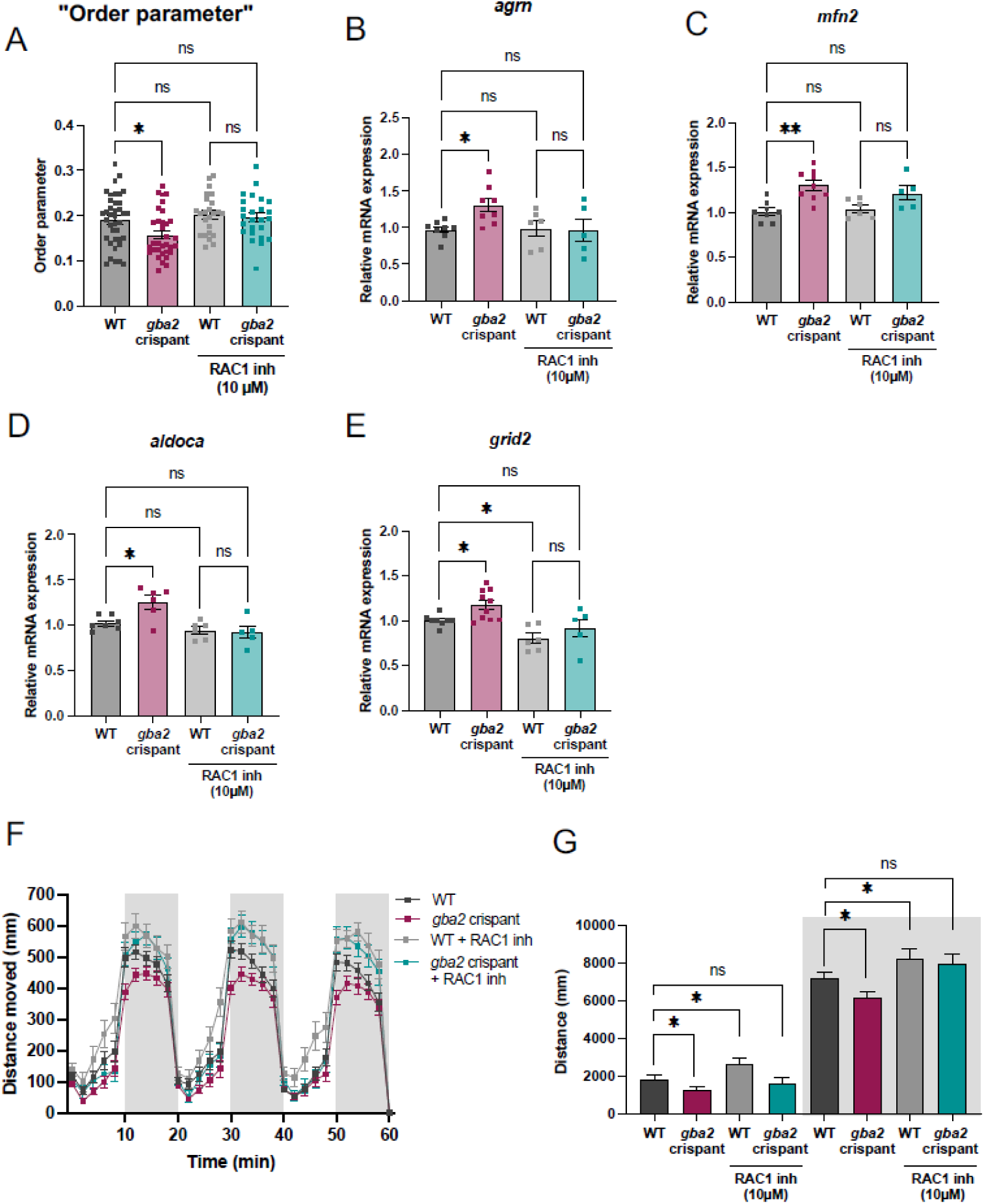
Treatment with NSC23766 Rac1 inhibitor partially rescues pathologic phenotypes in *gba2* crispant zebrafish larvae. (A) Quantification of “Order parameter” in 3-dpf WT and *gba2* crispants after 24h-treatment with NSC23766 (*n*≥24 from 3 independent experiments, Kruskal-Wallis test, WT vs *gba2* crispant, p=0.0343; WT vs WT+NSC23766, p=0.7526; WT vs *gba2* crispant+NSC23766, p=0.8875; WT+NSC23766. vs *gba2* crispant+NSC23766, p>0.8875). (B-C) Relative mRNA expression of cytoskeleton- and mitochondria-related genes in WT and *gba2* crispants after 24h-treatment with NSC23766 (*n≥*5 pools of 5-dpf zebrafish larvae from 3 independent experiments, Brown-Forsythe and Welch ANOVA tests for *agrn* (B), WT vs *gba2*, p=0.0453; WT vs WT+NSC23766, p>0.999; WT vs *gba2* + NSC23766, p>0.999; WT+NSC23766 vs *gba2* +NSC23766, p>0.999; and *mfn2* (C), WT vs *gba2*, p=0.0035; WT vs WT+NSC23766, p=0.9893; WT vs *gba2* +NSC23766, p=0.2423; WT+NSC23766 vs *gba2* +NSC23766, p=0.3908). (D-E) Relative mRNA expression of cerebellum-related genes in WT and *gba2* crispants after 24h-treatment with NSC23766 (*n≥*5 pools of 5-dpf zebrafish larvae from 3 independent experiments, one-way ANOVA for *aldoca* (D), WT vs *gba2*, p=0.0193; WT vs WT+NSC23766, p=0.6838; WT vs *gba2* +NSC23766, p=0.5323; WT+NSC23766 vs *gba2*+NSC23766, p=0.9905; and Brown-Forsythe and Welch ANOVA tests for *grid2* (E), WT vs *gba2*, p=0.04; WT vs WT +NSC23766, p=0.0477; WT vs *gba2* +NSC23766, p=0.8663; WT+NSC23766 vs *gba2* +NSC23766, p=0.7874). (F) Analysis of swimming behavior in 5-dpf WT and *gba2* crispants after 24h-treatment with NSC23766. Data are presented as the mean distance covered over time by *n*≥36 larvae from 3 independent experiments. (G) Quantification of the distance moved during the light and dark periods (*n*≥36 from 3 independent experiments, unpaired Student’s *t-*test for WT vs *gba2*, p=0.0375; WT vs WT+NSC23766, p=0.0456; WT vs *gba2* +NSC23766, p=0.5776 during light phase; WT vs *gba2*, p=0.0147, WT vs WT+NSC23766., p=0.0374, WT vs *gba2* +NSC23766, p=0.1305 during dark phase).

## Discussion

*GBA2*-associated HSP SPG46 is a rare neurodegenerative disorder associated with impaired lipid signatures for which no disease-modifying therapies exist. While GBA2 deficiency has been linked to cytoskeletal dysfunction and defective neurite outgrowth in immature neurons, the consequences on neuronal maturation and synaptic function, and the identity of a possible lipid mediator in the GBA2 catabolic pathway, remained unexplored. Here, we show that GlcChol depletion downstream of GBA2 inhibition drives a cascade of neuronal defects, i.e. presynaptic disorganization, NMDA receptor redistribution, altered Ca^2+^ signaling and mitochondrial dysfunction, converging on dysregulation of actin cytoskeleton via RAC1. Pharmacological inhibition of Rac1 rescued motor deficits and neuronal disorganization in a zebrafish *gba2* crispant model, suggesting that the GlcChol-RAC1-actin axis can serve as a tractable therapeutic target in SPG46.

### Stage-dependent neuronal defects persist functionally despite morphological compensation

Chronic GBA2 inhibition impairs axonal outgrowth in immature CGNs, consistent with previous reports [18], but morphological deficits resolve by DIV9, suggesting active compensatory remodeling. This morphological normalization, however, does not extend to synaptic function: at DIV14, structural and functional deficits persist, including presynaptic terminal enlargement, vesicle pool disorganization and divergent Ca^2+^ responses to depolarizing and glutamatergic stimuli. This dissociation underscores the importance of functional readouts in assessing disease-relevant pathology and suggests that compensatory mechanisms are insufficient to maintain synaptic integrity in mature neurons.

The divergent Ca^2+^ phenotype, i.e. reduced KCl-evoked and enhanced glutamate-evoked signals, points to dysfunction at distinct compartments. Reduced KCl-evoked responses are consistent with presynaptic structural alterations and dysregulated vesicle dynamics, whereas enhanced glutamate-evoked responses may reflect postsynaptic receptor dysfunction. These mechanisms are unlikely to be entirely independent, as depolarization-induced activation of VGCCs and glutamate-dependent activation of NMDARs represent interconnected sources of neuronal Ca²⁺ influx [38,39]. Pharmacological and imaging data further identify NMDAR redistribution as the primary driver of the enhanced glutamate response, independent of changes in total receptor levels or mitochondrial impairment. Exogenous GlcChol produced the opposite redistribution, directly implicating GlcChol homeostasis in the spatial regulation of NMDAR at the synapse, likely through RAC1-dependent actin reorganization [59,60]. Whether RAC1 directly mediates this redistribution downstream of GlcChol depletion remains to be demonstrated, but the possibility that alterations in cholesterol-derived lipid species may converge on NMDAR modulation through additional mechanisms, as suggested by the allosteric activity of 24(S)-hydroxycholesterol on NMDARs [61], warrants further investigation.

### Mitochondrial alterations are downstream consequences of cytoskeletal dysfunction

Despite model-specific differences in the molecular regulators involved, GBA2 deficiency consistently resulted in a mitochondrial remodeling phenotype across experimental systems. GBA2-inhibited CGNs showed enlarged mitochondria by TEM, accompanied by altered TMRM fluorescence and reduced MFN1 and TOM20 levels, suggesting changes in mitochondrial architecture and outer mitochondrial membrane organization. Similar alterations were observed in patient-derived fibroblasts carrying GBA2 loss-of-function mutations and in *gba2* crispant zebrafish, supporting the translational relevance of this phenotype. The increased TMRM signal likely reflects remodeling of the mitochondrial network rather than selective hyperpolarization, consistent with the enlarged mitochondrial morphology observed by TEM and with the proportional increase of both TMRM and MitoTracker signals in GBA2-HSP fibroblasts. Accordingly, these findings suggest that mitochondrial alterations are unlikely to represent the primary cause of Ca^2+^ dysregulation in this model. The link between RAC1 activity, actin dynamics and mitochondrial organization, motility and mitochondria-ER interactions [62,63] supports the interpretation that mitochondrial changes may arise from cytoskeletal dysfunction and/or altered GlcChol homeostasis rather than representing a primary pathogenic event. This contrasts with previous findings on truncated GBA2 mutants, which were reported to localize to the mitochondrial matrix and induce mitochondrial fragmentation and depolarization [64]. The opposite phenotype observed in our inhibition model and in GBA2^Cys688Thr^ patient fibroblasts suggests that mitochondrial consequences of GBA2 dysfunction critically depend on the molecular nature of the defect.

An additional mechanism potentially linking glucosylated sterol metabolism to mitochondrial remodeling involves neuronal HBB-B1. HBB-B1, which localizes to neuronal mitochondria and supports mitochondrial respiration [49], was reduced at the protein level in both GBA2-inhibited and GlcChol-treated CGNs. Since HBB-B1 depletion impairs neuronal oxygen homeostasis and has been associated with neurodegeneration [65], its reduction may contribute to the mitochondrial alterations observed in this model. Consistently, *Hbb* mRNA levels were reduced in zebrafish exposed to β-glucosyl sitosterol [15], supporting a role for glucosylated sterols in regulating HBB expression. Whether HBB downregulation directly contributes to mitochondrial remodeling or represents an adaptive response to altered lipid homeostasis remains to be established.

### Alterations in GlcChol level cause RAC1 dysregulation

Proteomic profiling of GBA2-inhibited and GlcChol-supplemented CGNs revealed a partial convergent signature of cytoskeletal and synaptic dysregulation, supporting the hypothesis that GlcChol should be maintained within a physiological range to support actin organization, and that deviation in either direction is detrimental. The reciprocal relationship between GlcChol levels and RAC1 abundance, increased upon GBA2 inhibition and reduced upon GlcChol supplementation, together with the reduction of active RAC1 and F-ACTIN signal observed upon GlcChol treatment, identifies RAC1 as a central mediator of cytoskeletal defects downstream of GlcChol dyshomeostasis. Consistently, alterations in actin cytoskeleton organization were also observed in patient-derived fibroblasts carrying the p.Cys688Thr GBA2 mutation, supporting the translational relevance of this axis. The causal role of RAC1 in the neuronal phenotype is further established by the rescue of motor deficits and neuronal disorganization in *gba2* crispant zebrafish upon pharmacological Rac1 inhibition, confirming that Rac1 dysregulation is not merely correlative, but functionally central to *GBA2*-HSP pathogenesis.

GBA2 inhibition is also associated with GlcCer accumulation, which could in principle contribute to the observed cytoskeletal phenotype. However, Raju *et al.* [17], while attributing the observed effects on RAC1 activity and cytoskeletal organization in *Gba2* KO and inhibited cells, did not provide direct evidence of GlcCer-mediated effects: all their data were obtained under conditions of global GBA2 loss or inhibition, where both GlcCer accumulation and GlcChol depletion co-occur. Similarly, cytoskeletal defects observed in *GBA1* mutant iPSC-derived neurons [66], where independent studies showed that GCase deficiencies also cause GlcCer accumulation but GlcChol increase [43,50], are consistent with a contribution of GlcChol dyshomeostasis alongside GlcCer accumulation. The finding that exogenous GlcChol supplementation alone recapitulates key proteomic and RAC1-mediated cytoskeletal alterations strongly implicates GlcChol as a primary lipid mediator, while a direct contribution of GlcCer accumulation cannot be formally excluded and warrants further investigation.

### Selective neuronal vulnerability reflects cell-type-specific dependence of GlcChol-RAC1 signaling relevant to HSP pathogenesis

The neuronal populations most affected in SPG46, corticospinal neurons, CGNs and Purkinje cells, may share characteristics that render them particularly sensitive to perturbations in RAC1-mediated actin dynamics. Corticospinal neurons depend on cytoskeletal integrity across extreme axonal lengths; CGNs operate high-frequency synapses with tight vesicle pool control; Purkinje cells require precise glutamate receptor organization at elaborate dendritic spines, a process highly cytoskeleton-dependent. Our *in vitro* findings in CGNs are complemented by evidence of cerebellar involvement in the *gba2* crispant zebrafish, where reduced cerebellar volume and altered *aldoca* and *grid2* expression suggest early Purkinje cell stress, analogous to the compensatory arborization preceding neurodegeneration in ataxic mouse models [67]. Whether this reflects cell-autonomous vulnerability to GlcChol-RAC1 dysregulation or secondary network effects remains to be determined, but the convergence of these observations on a single axis provides a possible mechanistic basis for the regional selectivity of SPG46 pathology. The co-occurrence of cytoskeletal, synaptic and mitochondrial defects in SPG46 resonates with emerging themes across HSP. Axonal cytoskeletal integrity is a shared vulnerability across multiple forms, including the most common, such as spastin (SPG4) and atlastin (SPG3A) [6,7,53]. The co-occurrence of cerebellar ataxia in SPG46 is part of a clinically recognized pattern: a subgroup of complex spastic ataxias, i.e. SPG35 (FA2H), SPG26 (B4GALNT1), SPG39 (PNPLA6), SPG5 (CYP7B1), SPG28 (DDHD1) and SPG56 (CYP2U1), is caused by mutations in lipid metabolism genes and consistently presents an ataxic component [53,68]. Our data place GBA2 and GlcChol at this intersection, suggesting that lipid-mediated RAC1 and actin cytoskeleton dysregulation may represent a convergent pathogenic mechanism across HSP forms with defective lipid metabolism.

### Rac1 as a therapeutic target

The *in vivo* rescue of SPG46 phenotypes by NSC23766 provides proof-of-concept evidence for RAC1 inhibition as a therapeutic strategy in this disease. NSC23766 was employed here as a mechanistic tool: its known pharmacological limitations are acknowledged [69], and more selective RAC1 inhibitors are available. One of them, JK-50561, tested in a phase II clinical trial for Alzheimer’s disease (ClinicalTrials.gov NCT05811442), supports the tractability of this target, even though results for the trial are still not available.

The therapeutic relevance of RAC1 is further supported by its emerging role across motor neuron diseases affecting overlapping neuronal populations. While GBA2-HSP involves RAC1 mislocalization and dysregulation, loss-of-function mutations in ALS2, encoding the RAC1 GEF ALSIN, cause juvenile ALS, primary lateral sclerosis and infantile-onset ascending HSP, all characterized by corticospinal neuron vulnerability, demonstrating that insufficient RAC1 activation is also pathogenic in this cell type [70,71]. Together, these observations suggest that RAC1 should be maintained within a homeostatic range, in terms of activity, abundance and localization, and that dysregulation in either direction is detrimental. This calls for careful mechanistic characterization before therapeutic translation, and future work in patient-derived iPSC neuronal models and stable *gba2* KO zebrafish lines will be essential to advance this strategy toward clinical application.

## Conclusions

This study identifies a pathogenic axis in SPG46 in which GBA2 loss of function leads to GlcChol depletion, RAC1 dysregulation and actin cytoskeleton disorganization, with downstream consequences on presynaptic organization, NMDA receptor distribution, Ca^2+^ signaling and mitochondrial dynamics. These defects are consistently observed across different *in vitro* and *in vivo* models, supporting their disease relevance. The pharmacological rescue of neuronal and motor phenotypes by RAC1 inhibition establishes this axis as a possible therapeutic target, and positions GlcChol homeostasis as a critical upstream regulator of neuronal function in the context of lipid-associated neurodegeneration.

## List of abbreviations

AchR: acetylcholine receptor
AGRN: agrin
ALDOCA: aldolase C
ALS: amyotrophic lateral sclerosis
ALSIN: amyotrophic lateral sclerosis 2 protein
AMP-DNM: AMP-deoxynojirimycin
B4GALNT1: beta-1,4-N-acetyl-galactosaminyltransferase 1
CGNs: cerebellar granule neurons
COFILIN: actin-depolymerizing factor (ADF/cofilin)
CYP2U1: cytochrome P450 family 2 subfamily U member 1
CYP7B1: cytochrome P450 family 7 subfamily B member 1
DDHD1: DDHD domain containing 1
DIV: days in vitro
DOK7: docking protein 7
DREBRIN: developmentally regulated brain protein
epo: erythropoietin
ER: endoplasmic reticulum
F-ACTIN: filamentous actin
FA2H: fatty acid 2-hydroxylase
FIS1: mitochondrial fission 1
FT: Fourier transform
GBA1: glucocerebrosidase 1
GBA2: non-lysosomal glucocerebrosidase 2
GCase: glucocerebrosidase
GD: Gaucher disease
GEF: guanine nucleotide exchange factor
GlcCer: glucosylceramides
GlcChol: β-D-glucosyl cholesterol
GRID2: glutamate receptor delta-2
HBB-B1: hemoglobin beta chain B1
HEK293: human embryonic kidney 293 cells
HIF1A: hypoxia-inducible factor 1-alpha
HOMER2: Homer scaffold protein 2
HSP: hereditary spastic paraplegia
iPSC: induced pluripotent stem cell
KCl: potassium chloride
KO: knockout
LRP4: low-density lipoprotein receptor-related protein 4
MFN1: mitofusin 1
MFN2: mitofusin 2
MUSK: muscle-specific kinase
NADH: nicotinamide adenine dinucleotide (reduced form)
NDUFAF1: NADH:ubiquinone oxidoreductase complex assembly factor 1
NMDAR: N-methyl-D-aspartate receptor
NMJ: neuromuscular junction
OPA1: optic atrophy 1
PAK1: p21-activated kinase 1
PD: Parkinson’s disease
PNPLA6: patatin-like phospholipase domain-containing protein 6
PSD: postsynaptic density
PSD-95: postsynaptic density protein 95
RAC1: Ras-related C3 botulinum toxin substrate 1
RAC1-GTP: GTP-bound active RAC1
ROS: reactive oxygen species
SOD1: superoxide dismutase 1
SOD2: superoxide dismutase 2
SPG: spastic paraplegia
TAGLN: transgelin
TEM: transmission electron microscopy
TMRM: tetramethylrhodamine methyl ester
TOM20: translocase of the outer mitochondrial membrane 20
VGCC: voltage-gated calcium channel
VGLUT1: vesicular glutamate transporter 1
WT: wild type
ZF: zebrafish.

## Declarations

**Ethics approval and consent to participate**

### Animal ethical statement

Our research complies with all relevant ethical regulations. Animal studies were conducted according to the Italian Ministry of Health and the approval by the Ethical Committee of the University of Padova and of the Italian Ministry of Health (authorization number 690/2020-PR). Experiments on zebrafish larvae were performed before the free-feeding stage and did not fall under animal experimentation laws according to EU Animal Protection Directive 2010/63/EU.

### Patient fibroblast ethical approval

The study was conducted in accordance with the Helsinki Declaration, and with the approval of the local ethics committee (IRB reference code CEPR 102/2020). Written informed consent was obtained from all subjects involved in the study.

## Consent for publication

Not applicable

## Availability of data and materials

The datasets used and/or analyzed during the current study are available from the corresponding author on reasonable request.

## Author contributions

A.C. performed the morphology and Ca^2+^ experiments on neurons. C.S. did the experiments on mitochondria and cytoskeleton in neurons and in patient fibroblasts. F.T. performed the zebrafish experiments. A.C, C.S. and F.T. prepared the figures and drafted the manuscript. L.P. contributed to the experiments performed on neuronal and zebrafish models. N.H. performed the proteomic experiments, analyzed the data and interpreted the results. S.M. adapted the use of AFT method to the zebrafish images and performed the analysis. N.F. and M.D.M. provided transgenic lines and contributed to the interpretation of the zebrafish work. L.L. supervised the proteomic experiments and provided expertise in sterol chemical biology. F.M.S. provided the patient fibroblast line and supervised the interpretation of the data in relationship with clinical information. L.D.V. conceived and supported the experiments on zebrafish models. N.P. conceived and lead the work, wrote and finalized the manuscript. All the authors revised and edited the final manuscript.

## Acknowledgements

We would like to thank Prof. Dagmar Wachten (University of Bonn, Germany) for providing the anti GBA2 antibody.

