## Supplementary figures for "A glucosylcholesterol-cytoskeleton axis links GBA2 loss-of-function to synaptic and mitochondrial pathology in Hereditary Spastic Paraplegia"

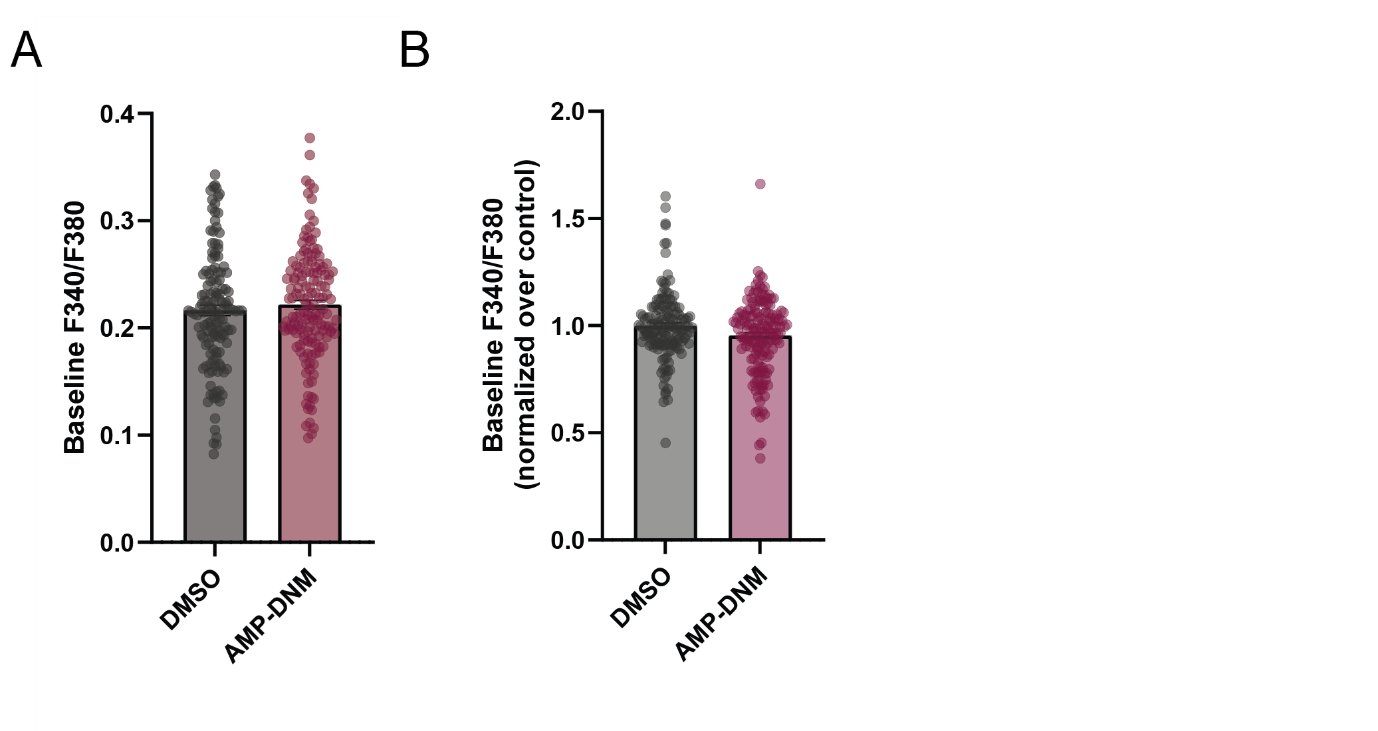

**Supplementary Figure 1: Basal calcium in GBA2-inhibited CGNs**

(A) Quantification of baseline Ca²⁺ prior KCl stimulation in CGNs (n ≥ 138 in 3 independent experiments, Shapiro–Wilk test for normality, Welch’s t test, p=0.4125). (B) Quantification of baseline Ca²⁺ prior glutamate stimulation (n ≥ 146 in 3 independent experiments, Shapiro–Wilk test for normality, Mann–Whitney test, p=0.14)

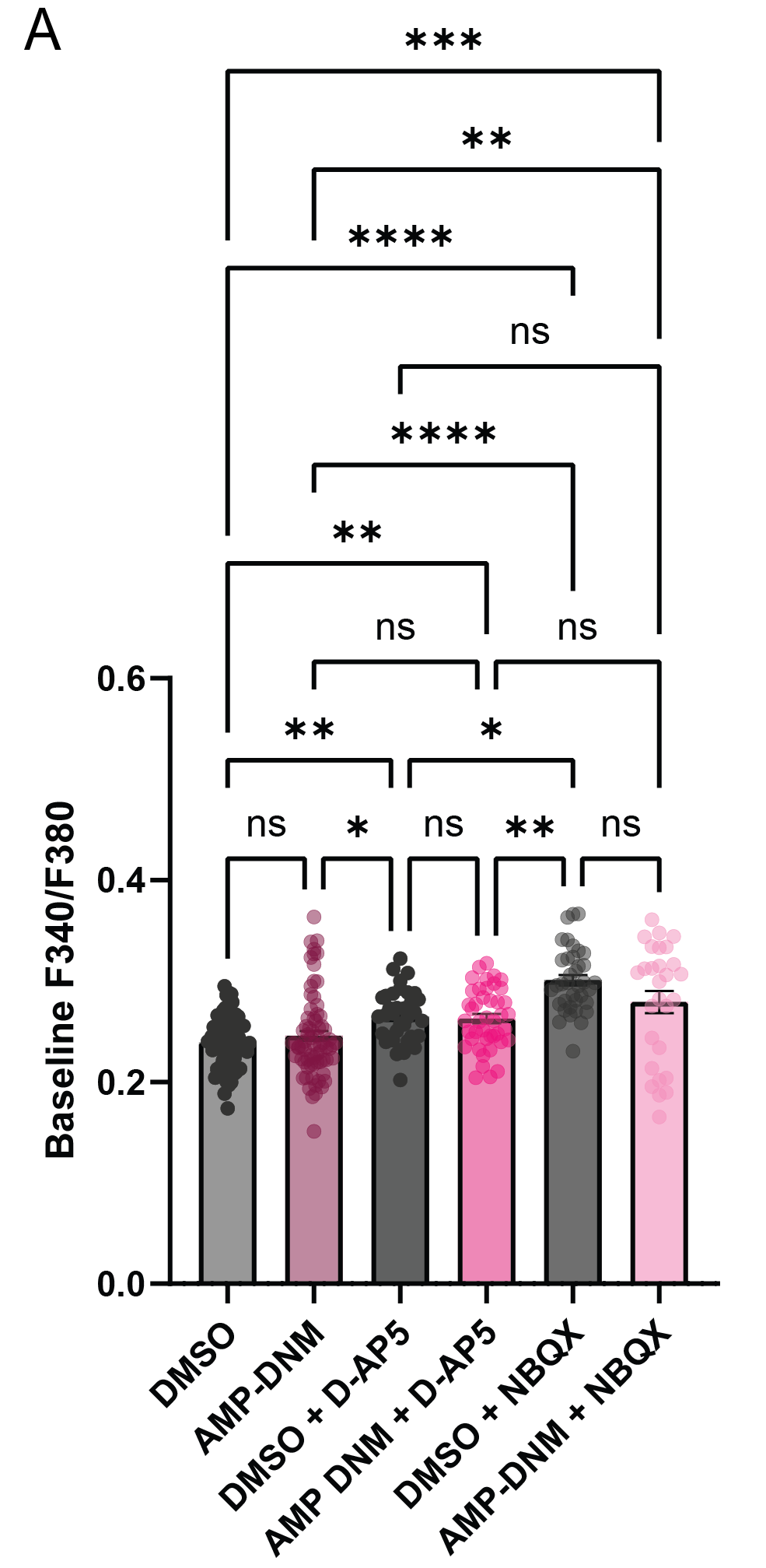

**Supplementary Figure 2: Basal calcium in GBA2-inhibited CGNs treated with D-AP5 (50 µM) or NBQX (25 µM)**

Quantification of baseline Ca²⁺ levels prior to glutamate stimulation in CGNs (n ≥ 26 cells from 3 independent experiments) in CGNs treated with D-AP5 (50 µM) or NBQX (25 µM). Shapiro–Wilk test for normality, Kruskal–Wallis test followed by Dunn’s multiple comparisons test (Adjusted p values for pairwise comparisons were as follows: DMSO vs. AMP-DNM, p > 0.9999 ; DMSO vs. DMSO + D-AP5, p = 0.0059; DMSO vs. AMP-DNM + D-AP5, p = 0.0085; DMSO vs. NBQX, p < 0.0001; DMSO vs. AMP-DNM + NBQX, p = 0.0004; AMP-DNM vs. DMSO + D-AP5, p = 0.0444; AMP-DNM vs. AMP-DNM + D-AP5, p = 0.0681); AMP-DNM vs. NBQX, p < 0.0001; AMP-DNM vs. AMP-DNM + NBQX, p = 0.0034; DMSO + D-AP5 vs. AMP-DNM + D-AP5, p > 0.9999; DMSO + D-AP5 vs. NBQX, p = 0.0126; DMSO + D-AP5 vs. AMP-DNM + NBQX, p > 0.9999; AMP-DNM + D-AP5 vs. NBQX, p = 0.0026; AMP-DNM + D-AP5 vs. AMP-DNM + NBQX, p > 0.9999; NBQX vs. AMP-DNM + NBQX, p = 0.3543.

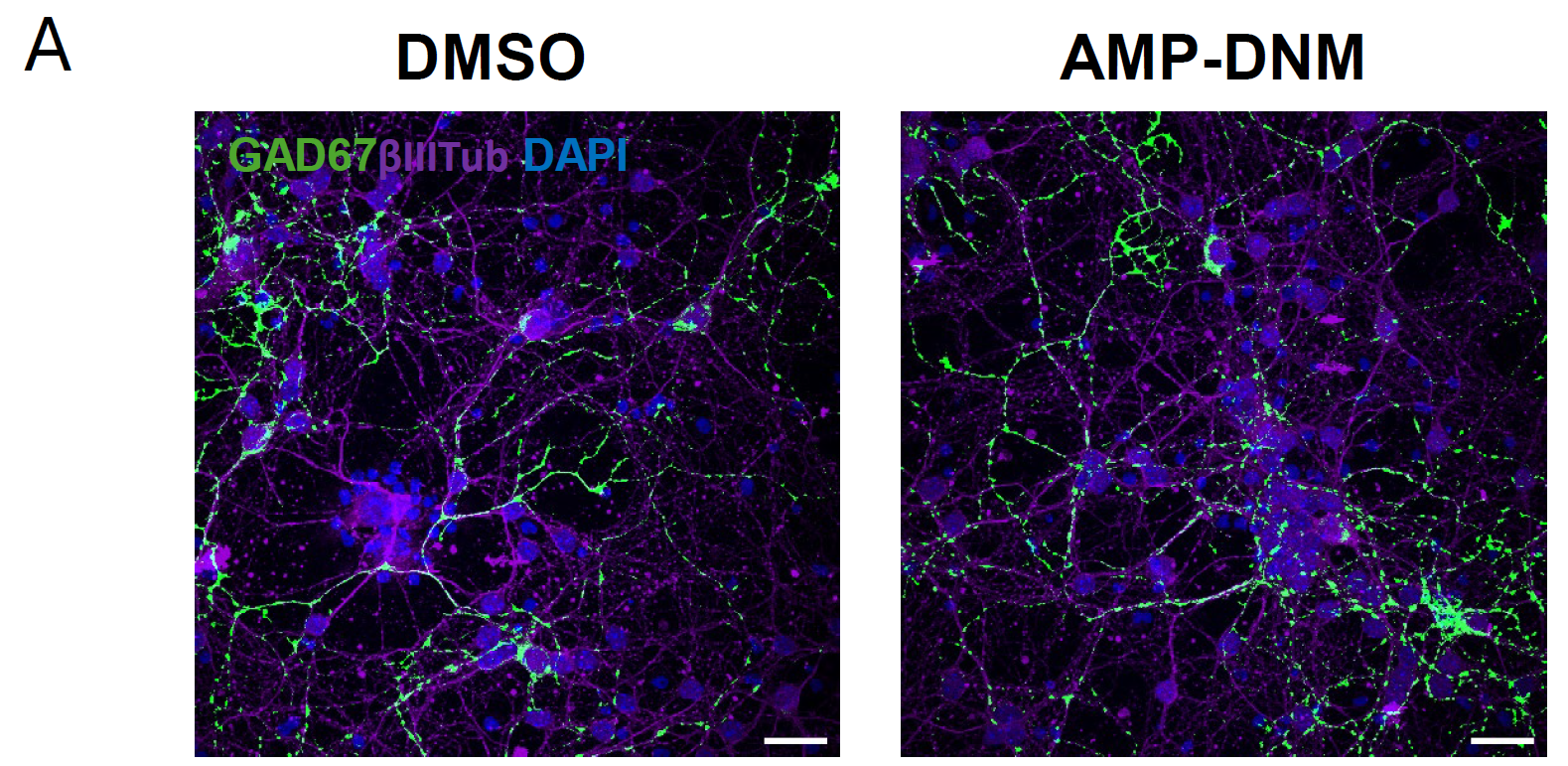

**Supplementary Figure 3: Visualization of inhibitory interneurons by GAD67 co-staining with βIII-tubulin**

Immunofluorescence staining of CGNs control (DMSO) and AMP-DNM-treated showing the overall neuronal network (βIII-tubulin-positive cells, neuronal marker, magenta) and the subset of GAD67-positive inhibitory interneurons. Nuclei were counterstained with DAPI (blue). (Scale bar: 20 µm).

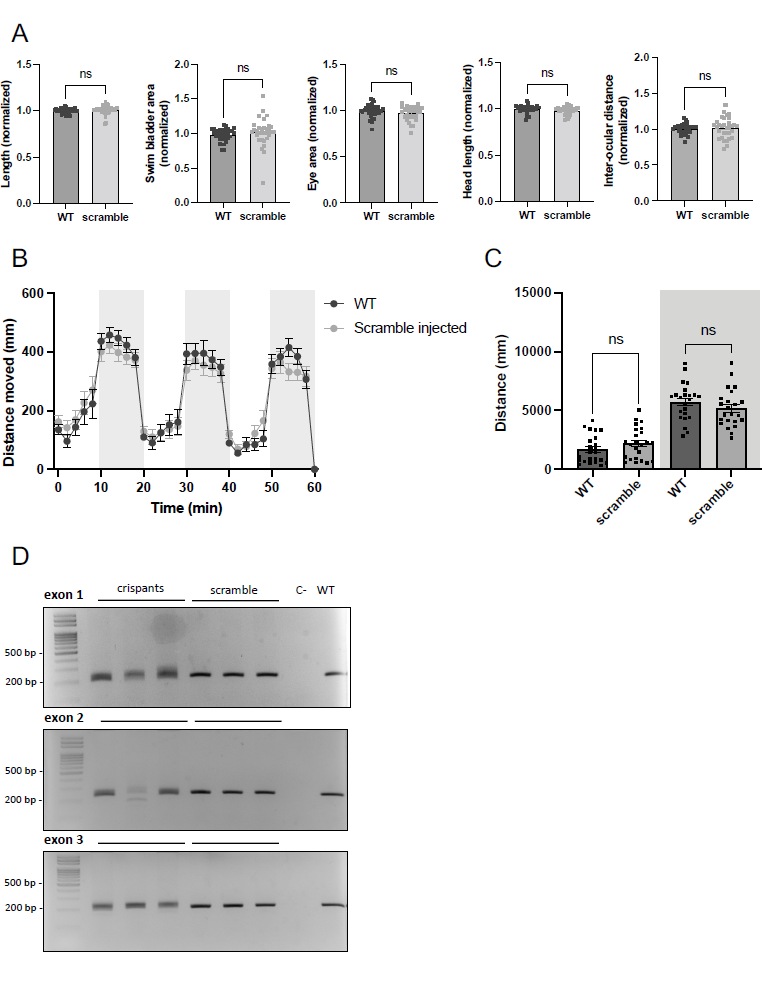

**Supplementary Figure 4: Morphometric, functional and molecular characterization of scramble injected zebrafish larvae.**

(A) Morphometric analysis of 5-dpf WT and scramble-injected zebrafish larvae (*n≥*32, Welch’s test for standard length, p=0.2837, swim bladder area, p=0.335, eye area, p=0.1451, head length/body length, p=0.1047 and interocular distance/head length, p=0.6597).

(B) Analysis of swimming behavior after alternating light/dark cycles stimulation. Data are presented as the mean distance covered over time by *n=*24 5-dpf zebrafish larvae.

(C) Quantification of the distance moved during the light (white background) and dark (grey background) periods (*n=*24 5-dpf larvae, Mann-Whitney test, p=0.1783 during light phase, p=0.1483 during dark phase).

(D) PCR analysis of genomic sequences targeted by gRNAs. Larval genomic DNA from injected *gba2* crispants (lanes 1-3) shows multiple bands due to Cas9 activity at the target sites in exons 1, 2, and 3 of the *gba2* gene. Genomic DNA from larvae injected with scrambled RNA (lanes 4-6) remains unaffected in the same regions. A non-injected WT larva (lane 8) was loaded as a reference.

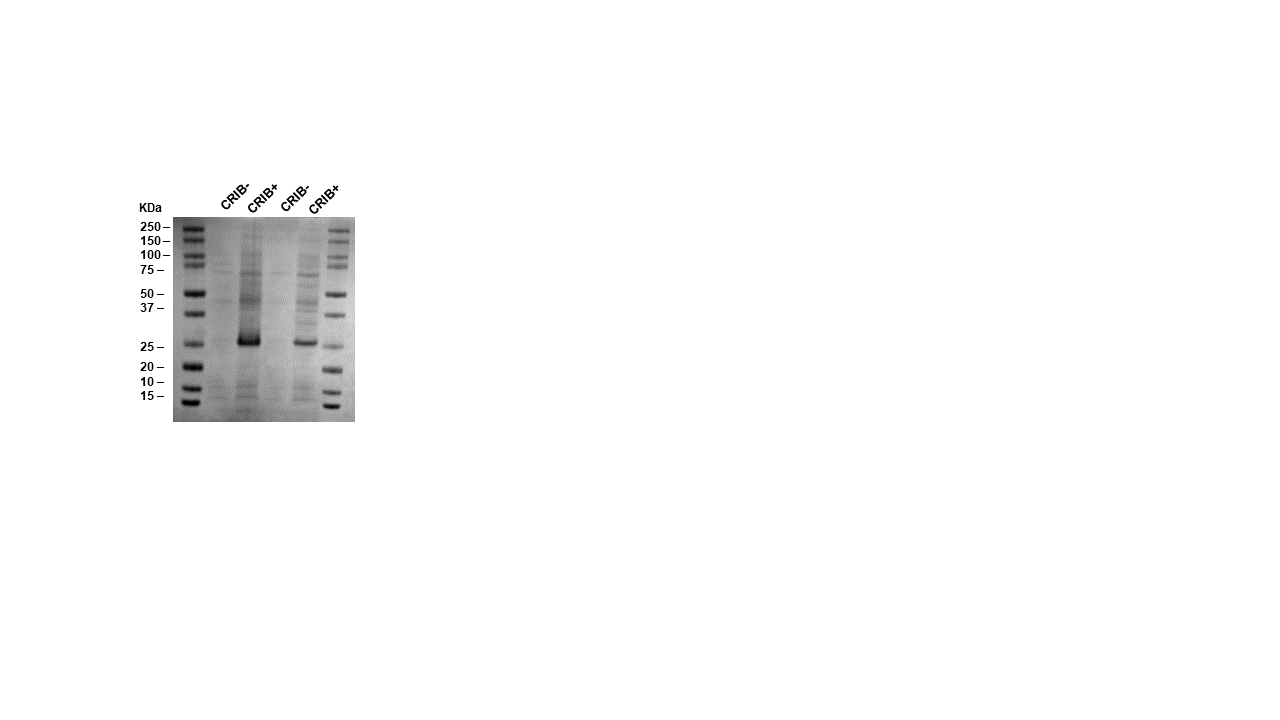

**Supplementary Figure 5: Validation of CRIB (Cdc42/Rac interactive binding)-PAK1 resin functionalization.** Coomassie-stained SDS-PAGE showing CRIB-PAK1-functionalized resin (CRIB+) and the corresponding non-functionalized resin control (CRIB-). The presence of the immobilized CRIB-PAK1 protein on the functionalized resin confirms successful protein coupling prior to Rac1-GTP pull-down assays.
