## Supplementary tables for "A glucosylcholesterol-cytoskeleton axis links GBA2 loss-of-function to synaptic and mitochondrial pathology in Hereditary Spastic Paraplegia"

**Table 1: RT-qPCR primers**

**Mouse primary cerebellar granule neurons**

| Gene | Forward sequence | Reverse sequence |
| --- | --- | --- |
| *Gba1* | TGCTAGTCGGTTCCAAGAGCTATGAT | TCTTTTCCCCCATCGATTTCAACCTT |
| *Gba2* | ACCAGACAGTCATTGCAGACCAATTTA | TGGTAGTCATGAGGCAAGATAGGTGT |
| *Gapdh* | AGGTCGGTGTGAACGGATTTG | TGTAGACCATGTAGTTGAGGTCA |

**Zebrafish**

| Gene | Forward sequence | Reverse sequence |
| --- | --- | --- |
| *gba1* | GCAGACTGATGGAGAGTCAAG | AATGTGCTGGAACTTCTGACTG |
| *gba2* | GTGTTGTACCACACGACATCG | GTACACCTGCAGCACAAACTTC |
| *agrn* | ACAGAGACGCATCAAGGTGATG | AGCTCAGAGGACACATACACTTG |
| *aldoca* | GGTATTACTATTGGGATCAAGGTTG | TACTGAGCACAGCGTTCTGAGA |
| *β-actin* | TGGGTATGGAATCTTGCGGT | GTGGGGCAATGATCTTGATC |
| *dok7* | CCTCCCTCCAACCGTCATC | TCTGTTCTCCATCTGCACACG |
| *epo* | GCCAGTATTAGAAACATCGCCA | CCCCGAAGAAAGTTGACATGG |
| *fis1* | AACTGGTTCACACAAGCAAAA | CCTGTTTGTTCCGGTTCA |
| *grid2* | CCGCATCGAGAACTCCATTC | CAGAGTCCAGAACCGTGC |
| *hif1α-3α* | AGAGCACACTTACCCACACA | ATGGGTGAGGTATGGGTTCG |
| *lrp4* | AGCAGGCTCTCAGCTGCTC | ACGGATGCTGGACAGGAGAC |
| *mfn2* | AACCGACGCAATGGATGTAC | CATCACGAATGGAGTCTGCC |
| *musk* | GACCAGCGCCTAAACCTGTC | ATGTGTTGAAAGCCACGCTGAG |
| *ndufaf1* | GGGACCTTGAGCTCCACC | GCCTCGTATCCGCAAATGTA |
| *opa1* | GGTACATCTTGCTTGGCTCAG | TTTCGCTCAGCTCCCAAAC |
| *sod1* | TCCGCACTTCAACCCTCATG | CATGAATCACCATGGTCCTCC |
| *sod2* | GGAGAGCGGAAGATTGAGG | TTGTACTGGAGATAGTACGCATG |
| *ef1a* | TTCGAGAAGGAAGCCGCTG | CAGCAACAATCAGCACAGCAC |

**Table 2: Zebrafish gRNAs**

| Gene | Gene direction  forward +, reverse - | crRNA’s spacer sequence | PAM sequence | Exon-target |
| --- | --- | --- | --- | --- |
| *gba2*-1EX | + | GGATTCGGTGTCCCCAAGGA | TGG | 1 |
| *gba2*-2EX | + | ATGGTGGTACCGAAAGACCC | AGG | 2 |
| *gba2*-3EX | - | CCTCCGCCAATGCCACCAAG | AGG | 3 |

**Table 3: PCR primers for *indel* formation analysis**

| Name | Forward sequence | Reverse sequence |
| --- | --- | --- |
| G-*gba2-*1EX | GTGCTGTTTTAGACTGGTGATC | GAAACATGCTCGAGCATGTTGG |
| G-*gba2*-2EX | CCTGTAATCCAACTTTATCCATCT | ATTTCAAGAGCTCTGATCACACTC |
| G-*gba2*-3EX | CACTGAGACAGATTTATGGTGAG | TAGCAGTCCATTACCTGATTAGC |
